# Human Neuron and Mouse Models Reveal Synaptic Imbalance in Kabuki Syndrome

**DOI:** 10.1101/2024.10.04.616738

**Authors:** Anna Wenninger, James Knopp, Steven Frye, Alexander Weaver, Neil McAdams, Sean Miller, Toshihiro Nomura, Ava Whitlark, Jack Swantkowski, Alexandra Noble, Caroline Lavender, Sandy Nam, Cameron MacKenzie, Isabella Wiebelt-Smith, Charles Sander, Katharyn Hutson, Jenny Bergqvist-Patzke, Sheri Sanders, Kasturi Haldar, Anis Contractor, Christopher Patzke

**Author notes:** Address correspondence to: Christopher Patzke, 109A Galvin Life Science Center, University of Notre Dame, Notre Dame, IN 46556. Contributed equally (Anna Wenninger, James Knopp).

## Abstract

Intellectual disability, affecting 2-3% of the general population, often co-occurs with neurodevelopmental disorders and is frequently caused by mutations that impair synaptic function. Kabuki syndrome (KS), a rare multisystem disorder associated with developmental delay and intellectual disability, results from mutations in either *KMT2D* (KS1) or *KDM6A* (KS2), encoding a histone methyltransferase and demethylase, respectively. The mechanisms underlying intellectual disability in KS remain poorly understood. Here, we generated human iPS cells carrying conditional Cre/lox-dependent loss-of-function mutations in *KMT2D* or *KDM6A* and differentiated them into excitatory or inhibitory neurons. Analysis revealed that KS1 and KS2 inhibitory neurons unexpectedly showed increased GABAergic synapse formation, whereas excitatory neurons displayed reduced synapse development and impaired synaptic transmission. We confirmed these findings in hippocampal neurons *in vitro* and *in vivo* using a mouse model for KS1 demonstrating a bidirectional shift: increase in inhibition and decrease in excitation. Synapse numbers and synaptic transmission in brain slices were shifted to increased inhibition/excitation ratio. Mechanistically, KS1 mutations activated neuroinflammatory signaling, impaired astrocytic function, and promoted glia-driven inhibitory synapse formation in human neurons. By integrating human neuron models with conditional *KMT2D*/*KDM6A* deletions and a *Kmt2d*-mutant mouse model, we identify a novel synaptic disease mechanism in KS that links chromatin remodeling defects to disrupted information transfer in neural circuits, providing a mechanistic explanation for intellectual disability.

**Significance Statement:** Kabuki syndrome, a rare disorder with intellectual disability, is caused by mutations in the chromatin regulators KMT2D and KDM6A, yet its cellular pathophysiology has remained unclear. Using new Cre/lox-inducible human neuron models together with a mouse model, we reveal a conserved excitation-inhibition imbalance characterized by increased inhibitory and decreased excitatory synapse formation. Our findings establish the first mechanistic framework linking *KMT2D*/*KDM6A* mutations to synaptic dysfunction, uncover glial contributions to disease pathogenesis, and suggest potential therapeutic avenues for restoring synaptic balance.

**Highlights:**

- KS1 and KS2 mutations cause decrease in excitatory synapses in human neurons
- KS1 and KS2 mutations cause increase in inhibitory synapses in human neurons
- KS1 mutation causes decrease in excitatory and increase in inhibitory synapses in mice
- KS1 mutant mouse astrocytes facilitate increase in inhibitory synapses in human neurons

## INTRODUCTION

Intellectual disability is a neurodevelopmental condition defined by significant limitations in intellectual functioning and adaptive behavior, often linked to abnormalities in synaptic structure and function ^1^. Kabuki syndrome (KS) is a rare multisystem disorder associated with developmental delay and intellectual disability, with an estimated prevalence of 1 in 32,000 births. It is caused by heterozygous/hemizygous loss-of-function mutations in either the *KMT2D* gene (KS1: 70% of cases) or the *KDM6A* gene (KS2: up to 5% of cases) ^2^. *KMT2D* (also called *MLL2* or *MLL4*) on chromosome 12q13 encodes a histone 3 lysine 4 methyltransferase (primarily H3K4me1), whereas the X-linked *KDM6A* (also called *UTX*) is a histone 3 lysine 27 trimethyl (H3K27me3) demethylase ^3–10^. Both proteins also act as tumor suppressors, with mutations reported in diverse cancers ^11–16^. KMT2D and KDM6A interact within the activating signal cointegrator-2-containing complex (ASCOM), which promotes open chromatin states at promoters and enhancers to regulate developmental gene expression ^7,17,18^. H3K4me1 marks active or primed enhancers and promoters, enabling enhancer–promoter interactions ^19^, and facilitating recruitment of histone H3K27 acetyltransferases (e.g., P300) and H3K27me3 demethylases (e.g., UTX) ^20^. KMT2D associates with chromatin remodelers at enhancers ^21^ and stabilizes KDM6A, consistent with reduced KDM6A levels in KMT2D-deficient cells, possibly explaining the broad overlap of clinical presentations^7^. Notably, *KDM6A* escapes X-inactivation and has a Y-chromosome homolog, *UTY*^22,23^, however, male KS2 patients present more severe manifestations than females ^24–26^.

KS1 and KS2 share core clinical features, including distinctive facial morphology (long palpebral fissures with eversion of the lower eyelid, resembling Kabuki theater makeup), postnatal growth deficiency, skeletal abnormalities, congenital malformations (e.g., cardiac and renal defects), immune dysregulation, and intellectual disability in up to 90% of affected individuals ^27–30^, occasionally without typical facial features.

Previous *in vitro* and animal studies have reported impaired hippocampal neurogenesis with associated memory deficits ^31–33^, defective differentiation of neural crest and progenitor cells ^34–37^, and preferential disruption of chromatin accessibility at CpG islands in mutant neurons ^38^. Yet, the mechanisms linking KS mutations to intellectual disability remain elusive. While extensive work has been done to identify and understand mutations contributing to intellectual disability ^1,39,40^, the specific effects in synaptic development and function underlying KS have not been defined.

Here, we investigate the impact of conditional KS1 and KS2 mutations on synaptic development and function in human excitatory and inhibitory neurons, complemented by a KS1 mouse model. We uncover a novel disease mechanism in which KS mutations impair excitatory synapse function while enhancing inhibitory synapse formation and transmission, leading to suppressed information flow and impaired network activity both *in vitro* and *in vivo*. Strikingly, neuroinflammation and astrocytic dysregulation promote excessive inhibitory synaptogenesis. Our findings, supported by morphological and functional analyses of synapses, electrophysiological and optical recordings of synaptic transmission, and assessments of neuronal and astrocytic activity, provide mechanistic insight into the basis of intellectual disability linked to KS.

## RESULTS

### Conditional heterozygous mutation in *KMT2D* increases synaptic inhibition and decreases excitation in human neurons

*KMT2D* is robustly expressed in wild-type human and mouse neurons (Suppl. Fig. 1). The human gene spans over 19 kilobases on chromosome 12 and contains 54 exons. To model KS1 in human neurons, we engineered a heterozygous conditional knockout in wild-type induced pluripotent stem cells (iPSCs) derived from a donor without disease-causing mutations, flanking exon 6 of *KMT2D* with loxP sites ^41,42^. Cre-mediated excision of exon 6 caused a frameshift, effectively creating a heterozygous knockout (Fig. 1A and 1B). We differentiated these engineered iPSCs into purely excitatory neurons by forced expression of the transcription factor Ngn2 ^43^, or purely inhibitory neurons by expression of the transcription factors Ascl1 and Dlx2 ^44^ (Fig. 1c and 1f). The mutation was induced via co-expression of Cre-recombinase. As a wild-type control, we used cells treated with an inactive mutant ΔCre (Fig. 1a). To validate haploinsufficiency we measured H3K4 mono-methylation in neurons and found a reduction in mutant cells (Fig. 1d and h), confirming decreased epigenetic activity in our model. Additionally, excitatory mutant neurons showed increased dendritic length and branching, but no change in axonal morphology (Suppl. Fig. 2). To examine synaptic morphology, we quantified puncta density, puncta area, and puncta intensity, which reflect synapse numbers and clustering. Inhibitory mutant human neurons exhibited increased vGAT puncta density compared to ΔCre controls, while excitatory mutant neurons showed decreased PSD95 puncta density (Fig. 1e, g, i and j) revealing opposite effects on GABAergic and glutamatergic synapses. We next assessed functional consequences. Using iGluSnFR3 imaging in excitatory neurons, we found reduced frequency of spontaneous glutamatergic events in mutants (Fig. 1h–k and Suppl. Fig. 3k and l), consistent with impaired release. Calcium imaging with GCaMP8m showed that excitatory mutant neurons fired fewer events per cell with larger amplitudes, while overall network synchrony remained unchanged (Suppl. Fig. 3a–e). Inhibitory neurons, in contrast, displayed reduced calcium activity both at the single-cell and average firing (purely inhibitory iNs don’t fire synchronously) (Suppl. Fig. 3f–j), possibly reflecting heightened inhibitory drive. Together, these results demonstrate that conditional heterozygous deletion of KMT2D induces morphological and functional synaptic changes, leading to impaired excitatory transmission and altered network activity across both excitatory and inhibitory neurons.

**Figure 1:**
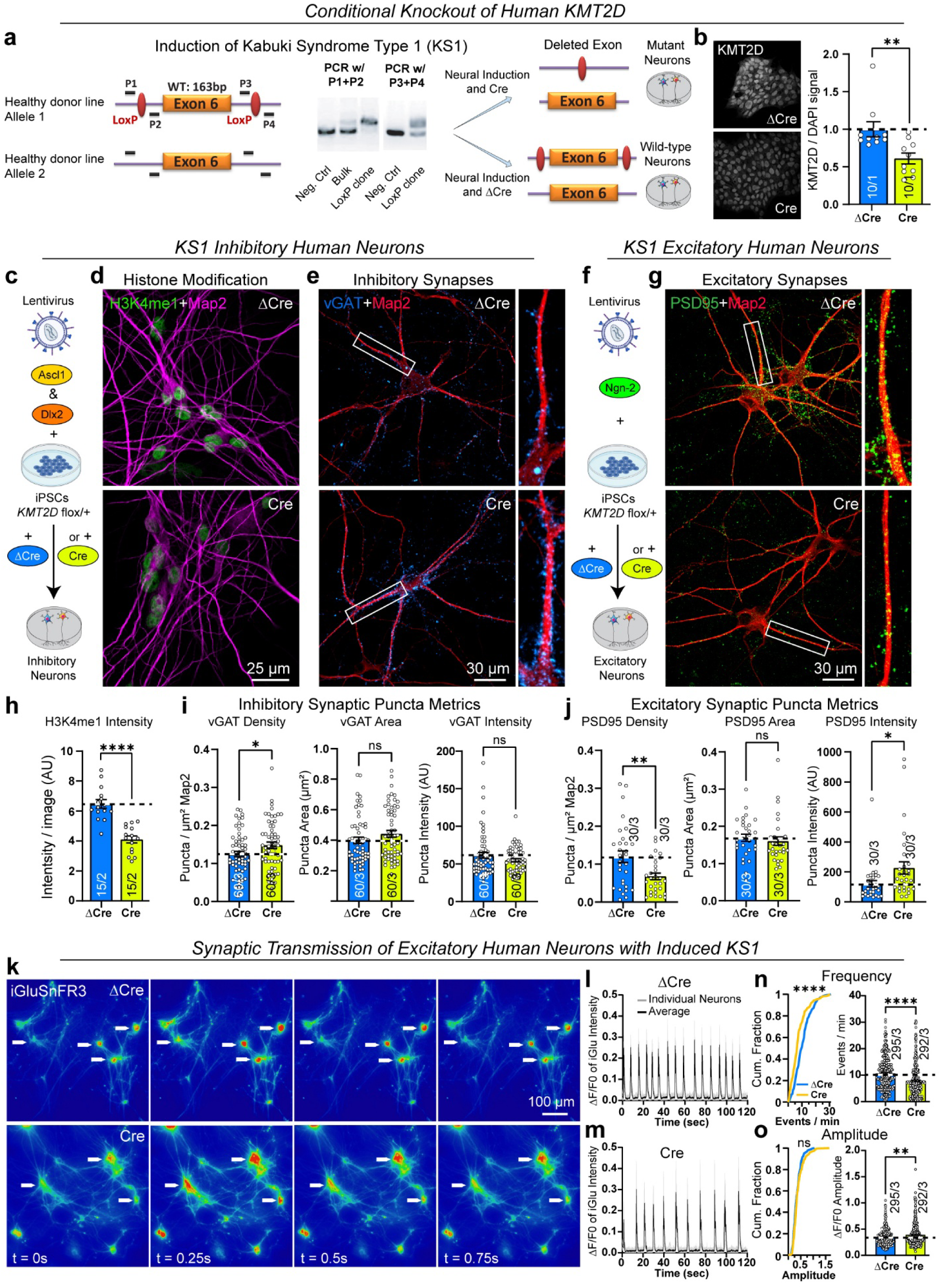
Increased synaptic inhibition and decreased excitation in a Cre/lox-dependent human model for Kabuki syndrome type 1 a, Strategy for Cre/lox-mediated induction of Kabuki syndrome type 1 (KS1) in human iPSCs from a donor without disease-causing mutations, and their differentiation into purely isogenic wild-type and mutant induced neurons (iNs). Specific heterozygous targeting of *KMT2D* and PCR-based screening strategy for correctly inserted loxP sites was performed by using primers P1+P2 for the upstream (333 vs 299bp, note that the upstream loxP site is inserted on both alleles) and P3+P4 for the downstream DNA sequences (302 vs 268bp). b, Immunofluorescent signal by anti KMT2D. Expression of active Cre resulted in ∼40% decrease of KMT2D signal as measured by fluorescent immunocytochemistry, compared to iPSCs treated with inactive ΔCre. c, d, e, h, and i, GABAergic (generated by lentiviral overexpression of transcription factors Ascl1 and Dlx2 in iPSCs(C)) *KMT2D*-mutant neurons (Cre) inhibitory neurons exhibit decreased Histone 3 lysine 4 mono-methylation (D and H) and an increased vGAT synaptic puncta density (e and i). f, g and j, Glutamatergic (generated by lentiviral overexpression of Ngn2 (f)) *KMT2D*-mutant (Cre) excitatory neurons show decreased PSD95 synaptic puncta density (g and j). k, l and m, Representative iGluSnFR3 micrographs of wild-type (ΔCre) and mutant (Cre) excitatory human neurons (10 frames apart) recorded at 40Hz frame rate and sample traces of fluorescence intensity changes (ΔF/F0). n and o, Left panels show cumulative distributions, and right panels display summary graphs of event frequency and amplitude for individual cells. Average values per field of view are shown in Suppl. Fig. 3k-l. Data shown are presented as individual values (each representing one image for confocal analyses or one cell for optical synaptic transmission recordings) with indicated mean ± SEM; number of cells or images analyzed and number of independent cell cultures are shown in graphs. Statistical significance was assessed with the Kolmogorov-Smirnov test for cumulative distributions and unpaired Student’s t test for summary graphs (*, p<0.05; **, p<0.01; **** p<0.0001).

### Conditional heterozygous mutation in *KDM6A* increases synaptic inhibition and decreases excitation in human neurons

We next asked whether KS2 mutations produce similar effects. KDM6A is strongly expressed in wild-type human and mouse neurons (Suppl. Fig. 4). To model KS2, we flanked exon 7 of *KDM6A* with loxP sites in wild-type iPSCs derived from the same male donor used for our KS1 model, thereby generating a conditional hemizygous knockout (Fig. 2a). Cre-expression led to efficient recombination and deletion of KDM6A signal in immunocytochemistry (Fig. 2a and b). Following differentiation into excitatory and inhibitory neurons, we observed increased vGAT puncta metrics in inhibitory neurons (Fig. 2c-e) and reduced PSD95 puncta metrics in excitatory neurons (Fig. 2f and g) without changes in dendritic or axonal arborization (Suppl. Fig. 5). Functional analysis with iGluSnFR3 imaging revealed reduced glutamatergic transmission in excitatory neurons (Fig. 2h–l and Suppl. Fig. 6g and h). GCaMP8m-based calcium imaging confirmed decreased activity in both individual cells and synchronously firing cells, alongside larger signal amplitudes in *KDM6A*-mutant neurons (Suppl. Fig. 6). Thus, the synaptic effects of *KDM6A*-mutant human neurons are similar to the phenotypes observed in *KMT2D*-mutant human neurons - increased inhibitory synaptic metrics and decreased excitatory neuronal communication.

**Figure 2:**
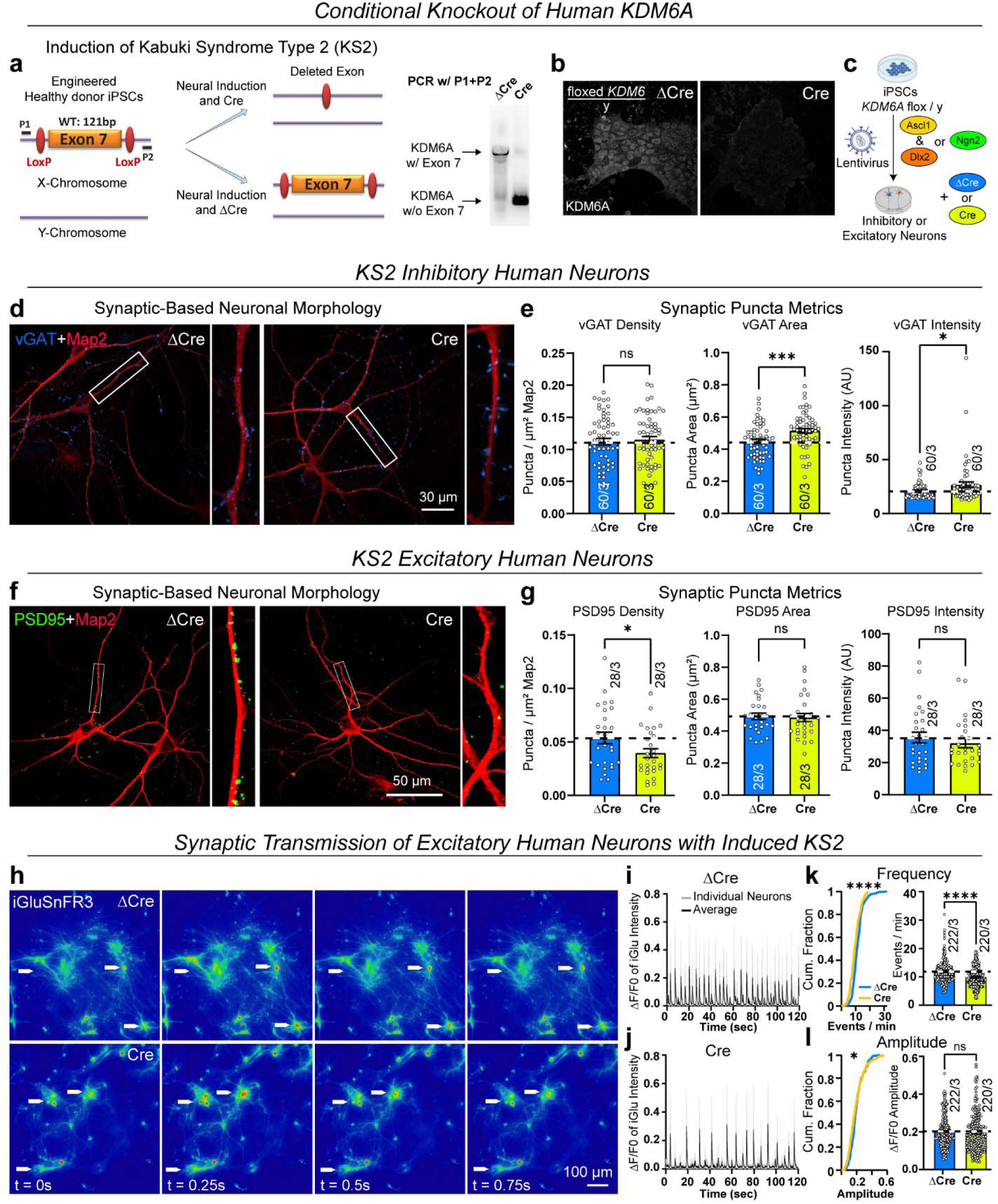
Increased synaptic inhibition and decreased excitation in a Cre/lox-dependent human model for Kabuki syndrome type 2 a, Hemizygous targeting strategy for Cre/lox-mediated induction of Kabuki syndrome type 2 (KS2) of X-linked *KDM6A* (also called *UTX*) in iPSCs from a male donor without disease-causing mutations and their differentiation into purely isogenic wild-type and mutant induced neurons. Correct targeting and successful recombination after expression of Cre was validated by PCR, using primers P1+P2 (ΔCre: 1219bp; Cre: 302bp). b, Expression of active Cre results in loss of KDM6A signal, as measured by fluorescent immunocytochemistry, compared to iPSCs treated with inactive ΔCre. c, Differentiation protocols used for generating GABAergic iNs (overexpression of transcription factors Ascl1 and Dlx2) of glutamatergic iNs (overexpression of Ngn2) for subsequent synapse analysis. d and e, *KDM6A*-mutant (Cre) inhibitory neurons show an increase in vGAT synaptic puncta area and intensity. f and g, *KDM6A*-mutant excitatory neurons show decreased PSD95 synaptic puncta density. h, i and j, Representative iGluSnFR3 micrographs of wild-type (ΔCre) and mutant (Cre) excitatory human neurons (10 frames apart) recorded at 40Hz frame rate and sample traces of fluorescence intensity changes (ΔF/F0). k and l, Left panels show cumulative distributions, and right panels display summary graphs of event frequency and amplitude for individual cells. Average values per field of view are shown in Suppl. Fig. 6g-h. Data shown are presented as individual values (each representing one image for confocal analyses or one cell for optical synaptic transmission recordings) with indicated mean ± SEM; number of cells or images analyzed and number of independent cell cultures are shown in graphs. Statistical significance was assessed with the Kolmogorov-Smirnov test for cumulative distributions and unpaired Student’s t test for summary graphs (*, p<0.05; ***, p<0.001; **** p<0.0001).

### Mouse model for Kabuki syndrome type 1 shows increased number of inhibitory and decreased number of excitatory synapses in hippocampal neurons

Next, we wanted to know if the KS1 mutation in mice showed similar effects as in KS1 and KS2 mutations in human neurons. We used heterozygous *Kmt2d*+/βGeo ^31^ animals to analyze synaptic morphology of hippocampal cells (Fig. 3a and 3b) ^45–47^. Immunolabeling for the pan-synaptic markers synapsin and bassoon revealed increased puncta metrics in mutant (Mut) compared to wild-type (WT) neurons (Fig. 3c-f), with the strongest increases proximal to the soma (Suppl. Fig. 7a-d). Dendritic length was also greater in mutants (Suppl. Fig. 7e–i), consistent with our findings in KS1 human neurons (Suppl. Fig. 2). To distinguish between excitatory and inhibitory changes, we labeled for PSD95 and vGluT1 (excitatory puncta metrics decreased, Fig. 3g-j) and for vGAT (same view field as vGluT1), Gephyrin and Neuroligin-2 (inhibitory puncta metrics increased, Fig. 3k-p), and confirmed the bidirectional synaptic shift: decreased excitation and increased inhibition. Because *Kmt2d*, is associated with defects in cell-type specific gene expression and cell differentiation, we next tested whether the number of neurons from mutant hippocampi was altered. Cell counts revealed fewer Map2/DAPI-positive neurons in mutant hippocampi, consistent with earlier studies ^31^, while glial numbers were unchanged (Suppl. Fig. 8a–c). Importantly, the number of inhibitory neurons (marked by somatostatin, parvalbumin, or Necab1) was unaltered (Suppl. Fig. 8d–g) ^48^. Additionally, we found a decrease in H3K4me1 signal when testing mutant neurons and glia cells, confirming the heterozygous epigenetic loss-of-function (Suppl. Fig. 8h and i). In summary, these data indicate that *Kmt2d*-mutant mouse hippocampal neurons exhibit a marked increase in inhibitory synapse metrics, accompanied by a decrease in excitatory parameters.

**Figure 3:**
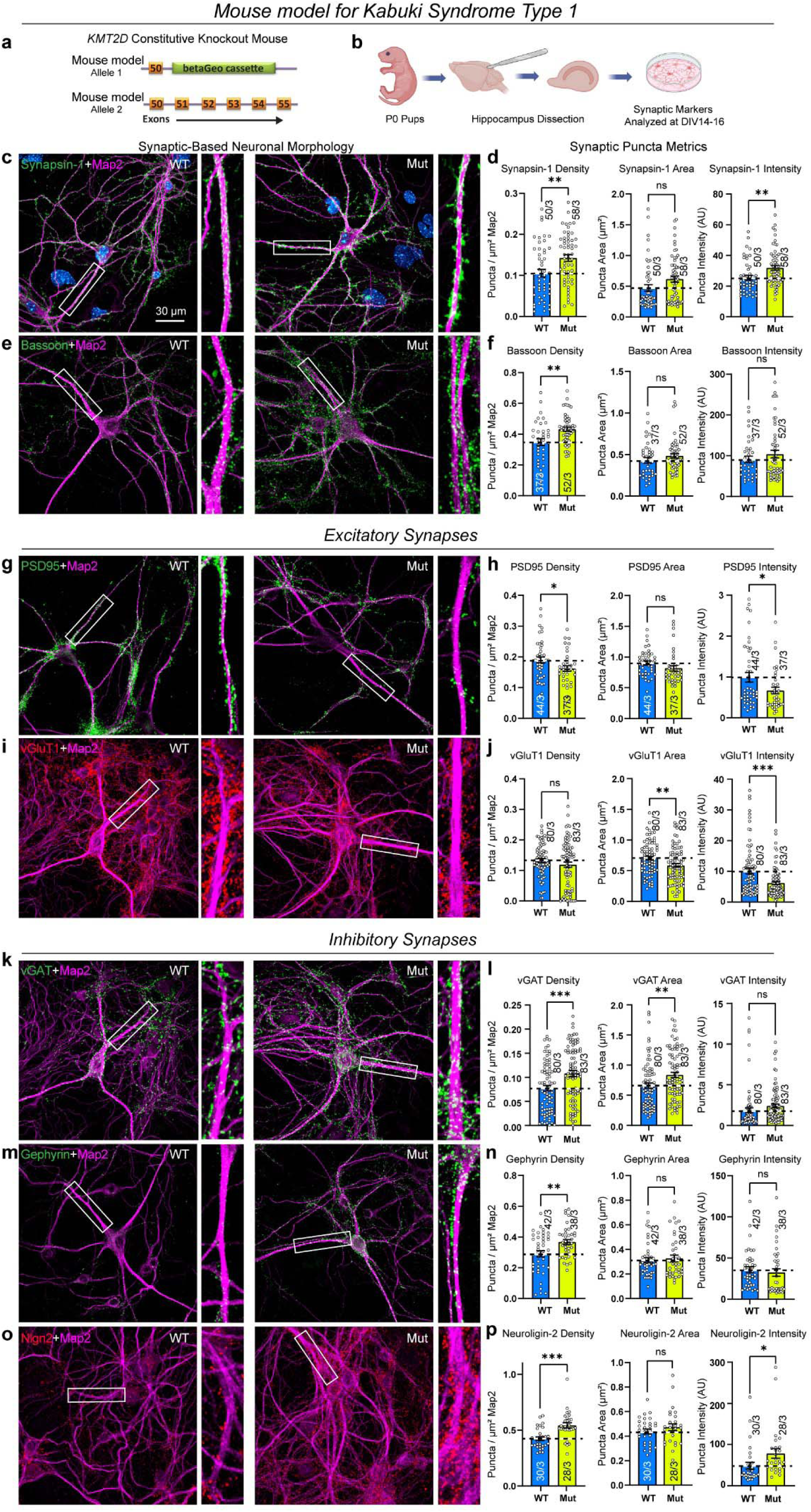
Decreased formation of excitatory and increased formation of inhibitory synapses in constitutive mouse model for Kabuki syndrome type 1 a and b, Schematic rationale of experiments: Newborn P0 mouse pups (littermates with (Mut) and without (WT) heterozygous mutation of *Kmt2d*) were used to analyze synaptic puncta metrics in cultured hippocampal cells. c-f, Pan-synaptic marker: Immunocytochemistry images showing synaptic morphology labeled for presynaptic Synapsin-1 (c) or Bassoon (e) (green), Map2 (magenta), and DAPI (blue). A zoomed-in view is included. Summary graphs show synaptic puncta metrics: density, puncta area and intensity. g-j, Excitatory synaptic markers: Immunocytochemistry images showing synaptic morphology labeled for postsynaptic excitatory PSD95 (g) (green) or presynaptic vGluT1 (i) (red), Map2 (magenta). A zoomed-in view is included. Summary graphs show synaptic puncta metrics. k-p, Inhibitory synaptic markers: Immunocytochemistry images showing synaptic morphology labeled for presynaptic inhibitory vGAT (k; same view field as i) (green), Gephyrin (m) (green) or Neuroligin-2 (o) (red), Map2 (magenta). A zoomed-in view is included. Summary graphs show synaptic puncta metrics. Data are presented as individual values (each representing one image) and means ± SEM; number of images analyzed and number of independent cell cultures are shown in graphs; statistical significance was assessed by two-tailed unpaired Student’s t-test (*, p<0.05; **, p<0.01; *** p<0.001).

### Bidirectional change of synaptic transmission in *Kmt2d*-mutant mouse hippocampal neurons

To evaluate functional consequences of the observed synaptic changes, we performed iGluSnFR3 and iGABASnFR2 imaging of hippocampal neurons *in vitro*. Mutant cells exhibited reduced excitatory and increased inhibitory neurotransmitter release, with corresponding changes in event frequency (Fig. 4 and Suppl. Fig. 9f–i), closely paralleling findings in KS1 human neurons (Fig. 1). Additionally, calcium imaging revealed reduced activity rates but larger amplitudes in mutant neurons (Suppl. Fig. 9). Next, we investigated synaptic effects *in vivo*: Hippocampal sections of P30 young adult mice showed increased inhibitory synapse formation in mutant animals (increased vGAT intensity, Fig. 5a-e and Suppl. Fig. 10) along with increased evoked inhibitory (IPSCs: inhibitory postsynaptic currents) and decreased excitatory synaptic transmission (EPSCs: excitatory postsynaptic currents) recorded from the same pyramidal neurons after stimulation of Schaffer collaterals in acute hippocampal slices (Suppl. Fig. 11), leading to a marked shift in the inhibition-to-excitation (I/E) ratio in KS1-mutant animals (Fig. 5f-h). In summary, these data demonstrate that the KS1 mutation in mice leads to increased inhibitory and decreased excitatory synapses within the hippocampus *in vitro* and *in vivo*.

**Figure 4:**
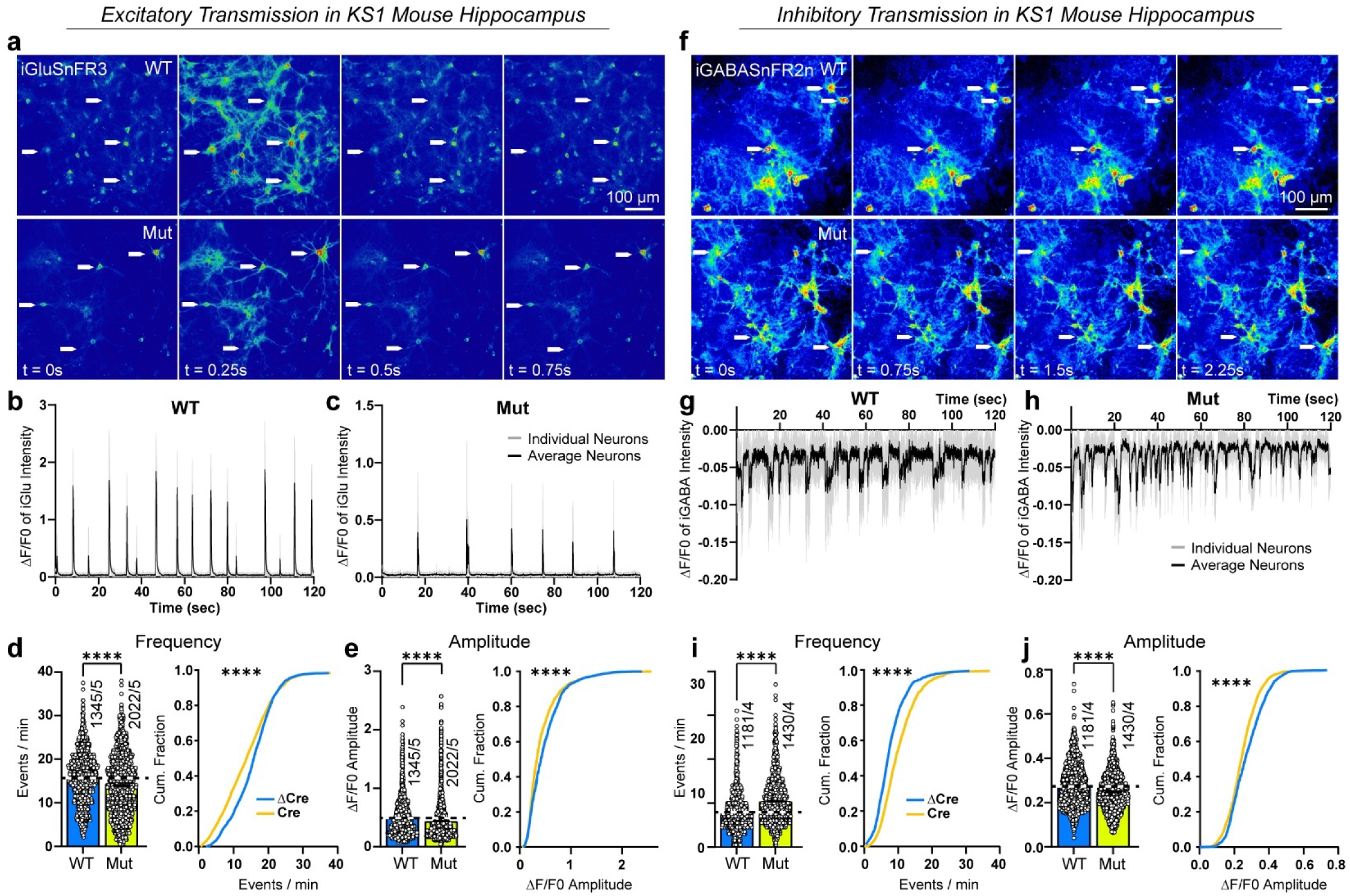
Bidirectional decreased excitatory and increased inhibitory synaptic transmission in constitutive mouse model for Kabuki syndrome type 1 a, Representative iGluSnFR3 micrographs of wild-type (WT) and mutant (Mut) cultured hippocampal cells (10 frames apart) recorded at 40Hz frame rate. b and c, Sample traces of fluorescence intensity changes (ΔF/F0). d and e, Left panels show summary graphs, and right panels display cumulative distributions of event frequency and amplitude. WT cells were derived from 8 animals and Mut cells from 20 animals across 5 litters. Average values per field of view are shown in Suppl. Fig. 9f-g. f, Representative iGABASnFR2 (negative-going reporter) micrographs of wild-type (WT) and mutant (Mut) cultured hippocampal cells (10 frames apart) recorded at 40Hz frame rate. g and h, Sample traces of fluorescence intensity changes (ΔF/F0). i and j, Left panels show summary graphs, and right panels display cumulative distributions of event frequency and amplitude. WT cells were derived from 9 animals and Mut cells from 16 animals across 4 litters. Average values per field of view are shown in Suppl. Fig. 9h-i. Data shown are presented as individual values (each representing one cell for optical synaptic transmission recordings) with indicated mean ± SEM; number of cells or images analyzed and number of independent cell cultures are shown in graphs. Statistical significance was assessed with the Kolmogorov-Smirnov test for cumulative distributions and unpaired Student’s t test for summary graphs (**** p<0.0001).

**Figure 5:**
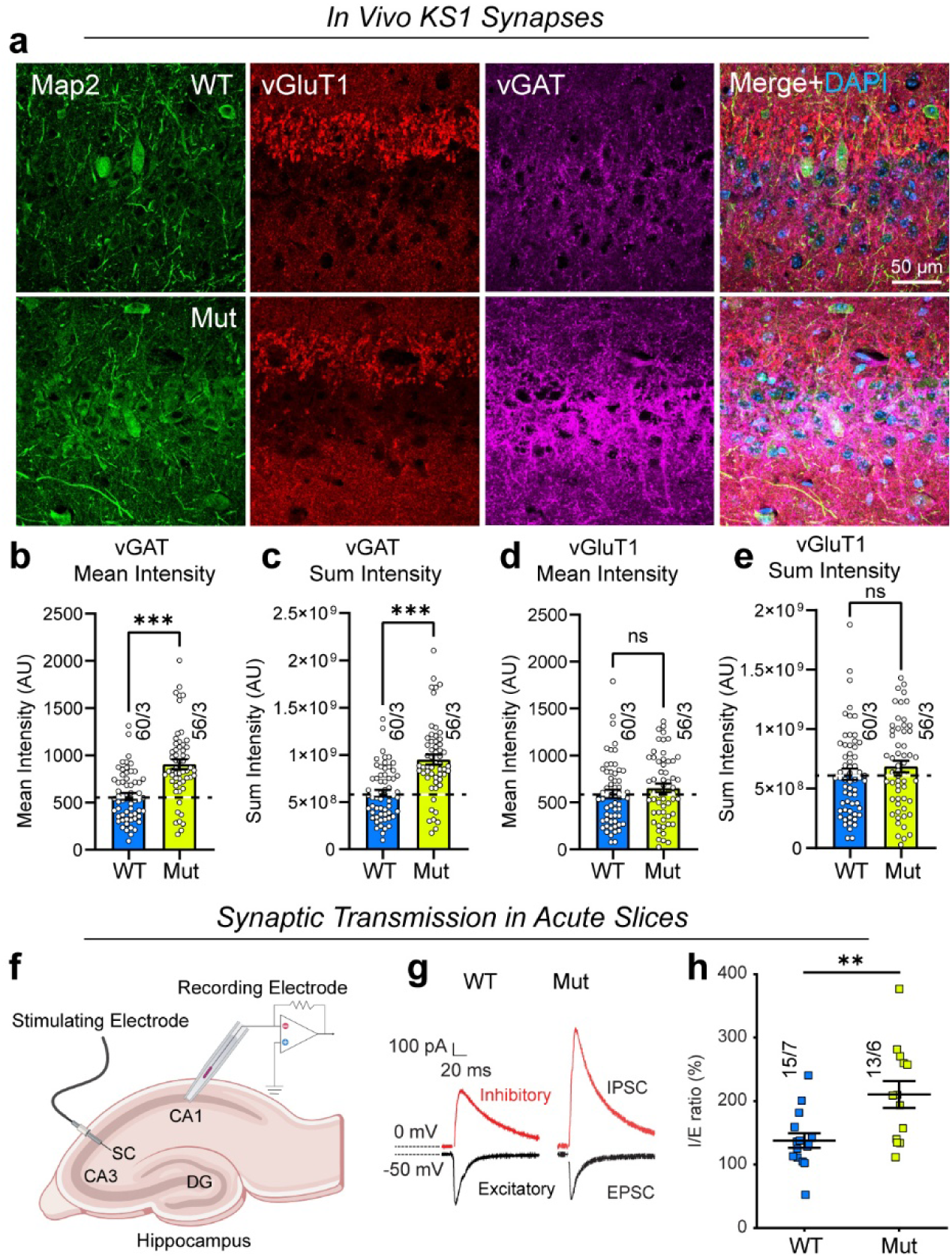
Increased number of functional inhibitory synapses in a constitutive mouse model for Kabuki syndrome type 1 *in vivo* a, Representative micrographs of hippocampal sections showing the CA3 region of WT and Mut mice. Sections are stained for Map2 (green), vGluT1 (red), vGAT (magenta), and DAPI (blue). b-e, Summary graphs for vGAT and vGluT1 staining showing mean intensity (intensity of all pixels/number of pixels) and the sum intensity of all pixels with the field of view. f, Experimental paradigm for recordings from acute hippocampal slices of WT and Mut mice. g and h, Representative traces of evoked IPSC (I) and EPSC (E) amplitudes and the summary graph showing their ratios. WT: 15 cells from 7 mice; Mut: 13 cells from 6 mice were recorded. Data are individual values (one image in b-e; and one cell in g, h) and means ± SEM; number of cells or images analyzed and number of independent animals are shown in graphs; statistical significance was assessed by unpaired Student’s t-test or Mann-Whitney U test (**, p<0.01; *** p<0.001).

### Neuroinflammation and astrocyte hypoactivity in *Kmt2d*-mutant mice reveal independent glial and neuronal contributions to excess inhibitory synapse formation

To investigate the underlying mechanism of the synaptic changes we performed RNA-sequencing on mouse hippocampal cells and detected 240 differentially expressed genes in the KS1 condition (Fig. 6a), with upregulation of immune response genes promoting neuroinflammation (*Cd36*, *Ccl12*, *Ccl9*, *Tgfb1*, *C3ar1*, *Iigp1c*, *Ifi211*) in combination with activation of immediate early genes (*Fos*, *Socs3*, *Atf3*, *Egr1*), changes linked to extracellular matrix remodeling and astrogliosis (*Fn1*, *Col8a1*, *Col5a2*, *Ltbp2*, *Thbs1*), and mixed regulation of genes affecting neuronal and synaptic development (with *Sema3b*, *Sema4g*, *Fgf11*, and *Sparcl1* downregulated and *Sema3f*, *Fst*, *Shh*, and *Igf1* upregulated in KS1 samples) (Fig. 6b and Suppl. Fig. 12). Interestingly, glutamate (*Slc7a11*, upregulated) and GABA (*Slc6a1* downregulated encoding GAT1, Suppl. Fig. 14a and b)) transporters were also differentially expressed suggesting altered extracellular neurotransmitter levels. Because many expression changes could be attributed to glia cells, which robustly express Kmt2d (Suppl. Fig. 12c, d, 13, 14a-c), we aimed to assess glial function in the presence of neurons via glia-specific recording of calcium activity using lentiviral mediated expression of F-GFAP-GCaMP8m in KS1 mouse hippocampal cultures. Interestingly, we saw a decrease in event frequency reflecting a decline in astrocytic activity (Fig. 6d-g). This prompted us to test how KS1-mutant mouse glia affect synapse formation in inhibitory human neurons. For this purpose, we generated inhibitory neurons (by overexpression of Ascl1 and Dlx2) with mutant (Cre) or wild-type (ΔCre) *KMT2D* (Fig. 1a) and co-cultured these with mouse glia with mutant (Mut) or wild-type (WT) *Kmt2d* (Fig. 3a and Fig. 6g). Strikingly, we detected an increase in inhibitory synaptic puncta metrics in KS1-mutant neurons co-cultured with wild-type glia but also in wild-type neurons co-cultured with KS1-mutant glia, suggesting independent contributions of mutant neurons and glia to increased inhibitory synapse formation (Fig. 6h and i, Suppl. Fig. 14d and e). Given that under the used conditions the vast majority of cultured glia cells are astrocytes, this hypoactive/reactive astrocyte state is sufficient to drive excessive inhibitory synapse formation via secreted or cell-surface molecules. These results indicate that complex transcriptional changes, including activated neuroinflammation, reduce astrocytic activity. Together with our neuronal data, this demonstrates a dual mechanism in KS1 pathogenesis: intrinsic neuronal defects combined with non-cell-autonomous contributions from reactive astrocytes.

**Figure 6:**
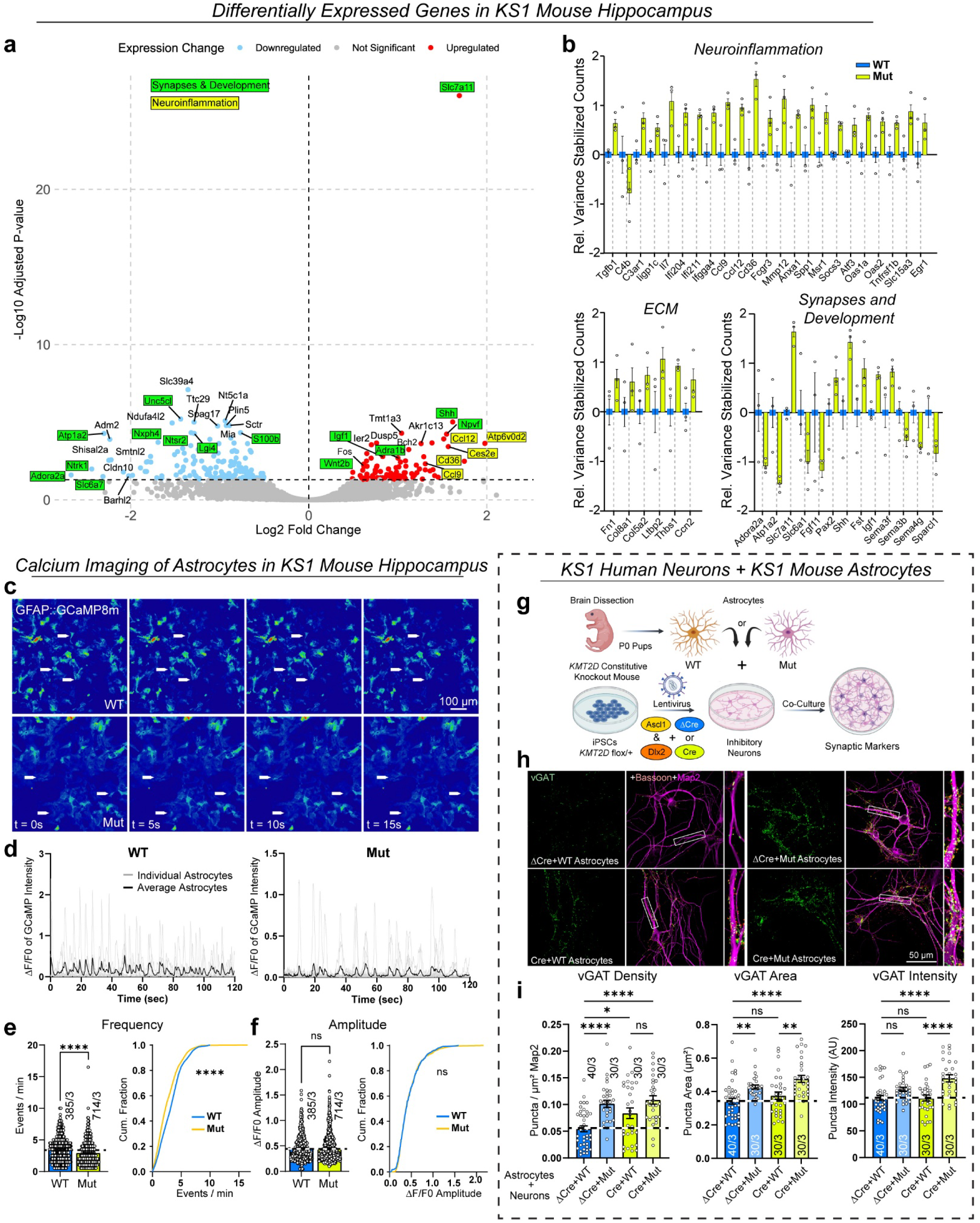
K*m*t2d haploinsufficiency (KS1) in the mouse hippocampus triggers glial neuroinflammation, driving increased inhibitory synapse formation in human neurons a, Volcano plot for mouse hippocampal cells showing up-and down-regulated in red and blue hues, respectively. Top differentially expressed genes are labeled, and highlighted in green or yellow for developmental or neuroinflammatory association, if applicable. Slc7a11 (also known as xCT) is a cystine/glutamate transporter. b, Variance stabilized expression values of select differentially expressed genes in KS1 mouse hippocampal cells (yellow) relative to wild type (blue). ECM: extracellular matrix. N=3 biological replicates from 3 experiments. c, Representative GFAP::GCaMP8m (expressed in astrocytes) micrographs of wild-type (WT) and mutant (Mut) cultured hippocampal cells (100 frames apart) recorded at 20Hz frame rate. d, Sample traces of fluorescence intensity changes (ΔF/F0). e and f, Left panels show summary graphs and right panels display cumulative distributions of event frequency and amplitude. g, Co-culture of human wild-type (ΔCre) or *KMT2D*-mutant induced inhibitory neurons (Cre) with wild-type (WT) or *Kmt2d*-mutant (Mut) mouse astrocytes. h and i, KS1 mutations in human neurons or mouse astrocytes are sufficient to increase inhibitory synaptic puncta metrics in human neurons. Data shown are presented as individual values (each representing one image for confocal analyses or one cell for astrocytic GCamP8m recordings) with indicated mean ± SEM; number of cells or images analyzed and number of independent cell cultures are shown in graphs. Statistical significance was assessed with the Kolmogorov-Smirnov test for cumulative distributions and unpaired Student’s t test for summary graphs in e and f; and one way ANOVA and Tukey’s post hoc test for multiple comparisons in i (*, p<0.05; **, p<0.01; **** p<0.0001).

## DISCUSSION

In this study we created and analyzed new Cre/lox-inducible human neuron model systems for KS1 (Fig. 1a) and KS2 (Fig. 2a) and complemented our findings with KS1 mice (Fig. 3a). We focused on the formation and function of synapses to gain mechanistic insights into the neurodevelopmental alterations leading to intellectual disability in KS patients – a link that had not previously been established. Aberrant synapse formation and function are recognized key drivers of neurodevelopmental disorders with intellectual disability ^49–52^. Across all three models, we found that heterozygous/hemizygous loss of KMT2D (or Kdm2d in mice) or KDM6A leads to increased inhibitory synapse formation and decreased excitatory synapse formation. These changes alter both synapse numbers and synaptic transmission in a bidirectional manner, producing an imbalanced inhibition-to-excitation ratio that dampens information flow and coordinated network activity (Fig. 7). This conclusion is supported by morphological and functional evidence: altered synapse numbers, impaired excitatory and inhibitory transmission, and disrupted neuronal calcium signaling. Furthermore, in KS1 mice we observed neuroinflammation with astrocytes displaying partial reactivity (modest GFAP elevation) and reduced calcium activity. This dual state may reflect secondary consequences of reduced neuronal activity and/or impaired neuron–glia communication at tripartite synapses. Importantly, mutant astrocytes were sufficient to drive increased inhibitory synapse formation in wild-type human neurons, while mutant human neurons could also induce this phenotype on wild-type astrocytes. Since all KS1 and KS2 human neuron experiments were performed in co-culture with wild-type mouse astrocytes by default, our findings suggest that inhibitory synapse formation is promoted by both neuronal intrinsic and astrocytic factors. Candidate glial drivers include neuroinflammatory genes. Additionally, we detected increased expression of Neuroligin-2, a postsynaptic cell adhesion molecule specific for inhibitory synapses. Given that Neuroligin-2 overexpression increases inhibitory synapse formation while deletion reduces inhibitory input ^53–55^, it likely contributes directly to the phenotype in our models. Increased inhibitory synapse formation cannot be explained by changes in neuron numbers. In KS1 mouse hippocampus, we observed more inhibitory synapses without an increase in inhibitory neuron counts, whereas the total number of neurons was reduced (Suppl. Fig. 8a–g), indirectly pointing to loss of excitatory neurons. More neurons would create more synapses. But here, fewer neurons create more synapses (as shown by the pan-synaptic markers Synapsin-1 and Bassoon (Fig. 3c-f)), with excessively formed inhibitory synapses outweighing the lost excitatory synapses. This finding of increased synapse formation (vs. increased number of inhibitory cells) also applies to human neurons, which are produced by transducing and plating the same number of iPSCs per genetic condition (Cre vs. ΔCre) in a standardized manner, leading to the same number of neurons. The generated human inhibitory neurons express mostly calretinin or calbindin ^56^ and represent distinct non-overlapping populations of interneurons found in the cerebral cortex (layers I-II and V) and hippocampus (dentate gyrus, the stratum moleculare and stratum lacunosum of the CA1 region). Together, this indicates that the increase in inhibitory synapses reflects an increase of synapse formation per cell and is not the result of increased neuron numbers.

**Figure 7:**
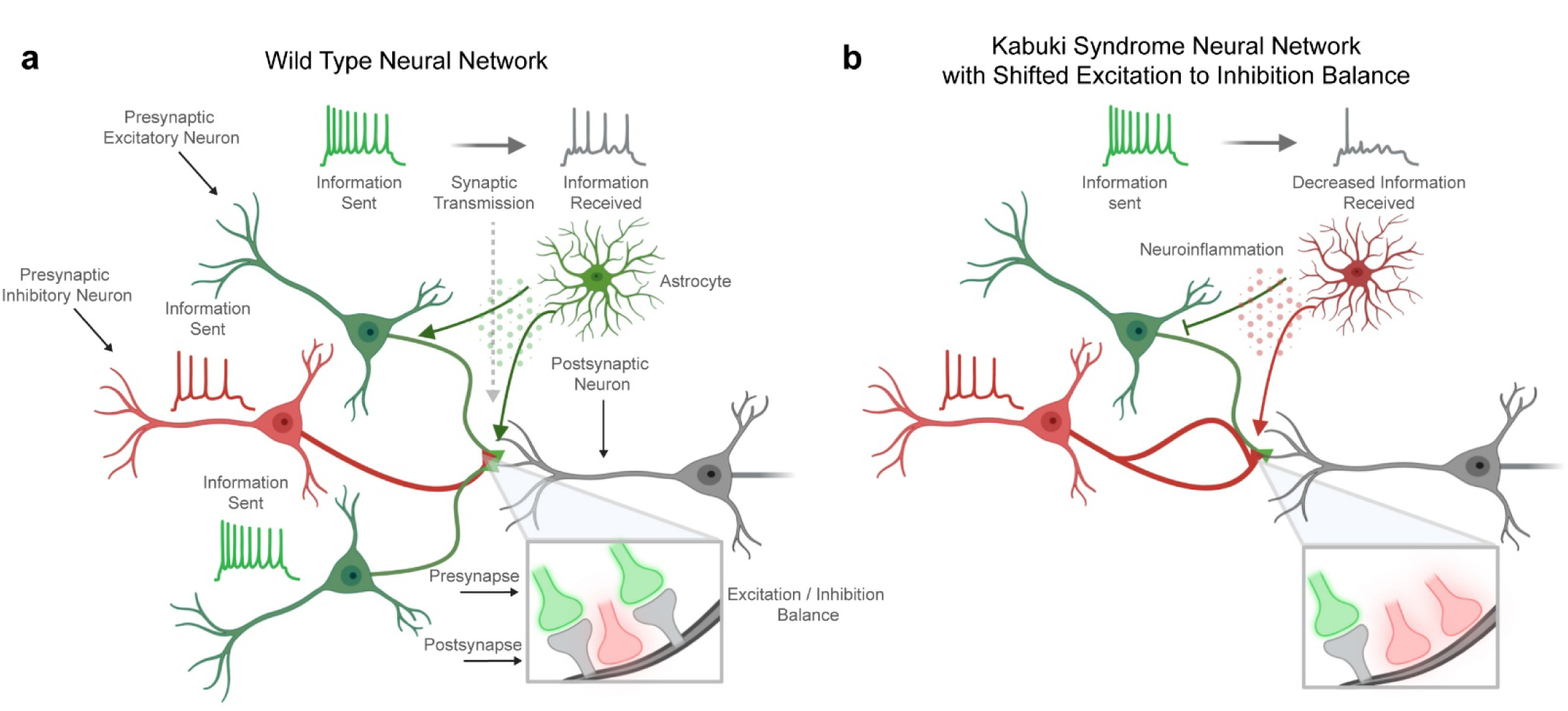
Bidirectional synaptic imbalance in Kabuki syndrome: decreased excitatory and increased inhibitory synapse formation. a, Example network consisting of excitatory (green) and inhibitory neurons (red), promoting and repressing information transfer at synapses (spike diagrams, left vs. right, respectively). b, Synaptic dysfunction: Due to an overabundance of inhibitory synapses, information transfer through the network is impaired. Distinct contributions of neurons and astrocytes are sufficient to induce increased inhibitory synapse formation.

Regarding the loss of excitatory synapses, the situation is different. It has been shown that Kmt2d deficiency in mice results in reduced neurogenesis in the hippocampus paired with memory deficits ^31^. Interestingly, the most significantly upregulated gene in our study is *Slc7a11*, which is also known as xCT – a cystine-glutamate antiporter, that brings extracellular cystine into the cell in exchange for intracellular glutamate, thereby controlling extracellular glutamate levels ^57^. In glioma, *Slc7a11* upregulation causes excessive glutamate release and excitotoxic neuronal death ^58^. Here, overexpression of *Slc7a11* (mainly expressed in astrocytes) in KS1 hippocampi could thus explain the observed partial loss of neurons: excitotoxicity leads to fewer excitatory neurons, which generate fewer excitatory synapses. Interestingly, many human cancers overexpress *Slc7a11*, which protects cells from oxidative stress–induced death (e.g., ferroptosis). Notably, epidermis-specific knockout of *Kmt2d*, used to model keratinocyte cancers, also causes Slc7a11 upregulation ^14,59^, consistent with our findings. Moreover, knockdown of elevated *Slc7a11* expression in the BTBR mouse model of autism spectrum disorder (ASD) was shown to reduce ASD-associated behavioral phenotypes ^60^. Strikingly, mice lacking cystine transport via xCT (*Slc7a11*) exhibit increased cell proliferation in neurogenic regions of the subventricular zone (SVZ) and dentate gyrus (DG) in the adult brain ^61^. This suggests that inhibiting Slc7a11 may bypass the reduction of hippocampal neurogenesis in KS1 mice, offering a potential therapeutic strategy for Kabuki syndrome type 1.

Additionally, we observed downregulation of *Sparcl1* (Hevin), encoding an astrocyte-secreted synaptogenic protein that bridges presynaptic neurexin-1α and postsynaptic neuroligin-1B (thereby promoting excitatory synapse formation), may also contribute ^62^. Reduced *SPARCL1* expression has been reported in neurological disorders such as autism and depression, and genetic alterations to *SPARCL1* (including copy number variations, polymorphisms, and mutations) are linked to autism and schizophrenia ^63–65^. An additional neuronal intrinsic factor explaining reduced excitatory synapse formation could be the upregulation of *Sema3f*, which has been shown to inhibit excitatory synapse formation by constraining the density of dendritic spines along apical dendrites of deep layer cortical pyramidal neurons in mice ^66^.

The molecular basis of these phenotypes likely resides in the chromatin remodeling functions of KMT2D and KDM6A. *KMT2D* mutations disrupt histone methylation, altering transcriptional programs ^12,67,68^. Our findings suggest that loss of function in either *KMT2D* or *KDM6A* represses gene networks required to maintain excitatory and inhibitory synaptic balance. Therapeutically, restoring this balance represents a promising strategy. In particular, pharmacological interventions that increase excitatory synaptic activity or decrease excessive inhibitory synaptic activity could help to rescue synaptic homeostasis. Several drugs which have been shown to enhance AMPA (Aniracetam, Piracetam) and NMDA (D-cycloserine) receptor activity as well as those shown to decrease GABA-A (Bicuculline, Gabazine) or GABA-B (CGP55845) receptor activity would serve as initial tools for studies involving the pharmacological modulation of I/E imbalance and for determining potential targets in treating Kabuki syndrome. Interestingly, previous research on the KS-related disorder Rubinstein-Taybi syndrome has uncovered evidence of altered neural development and consequential synaptic plasticity deficits ^69–71^. In conclusion, our study establishes I/E imbalance as a unifying pathophysiological mechanism across Kabuki syndrome models. By uncovering both neuronal and astrocytic contributions, we provide a framework for future mechanistic studies and therapeutic development aimed at restoring synaptic balance in this rare but informative neurodevelopmental disorder. At the same time, our results suggest that inhibiting KMT2D and KDM6A may promote inhibitory synapse formation.

## MATERIALS AND METHODS

### Mice

Animal experiments conducted in this study were in accordance with National Institute of Health Guidelines for the Care and Use of Laboratory Mice and approved by the Institutional Animal Care and Use Committee (IACUC) at the University of Notre Dame. The <u>Kmt2d+/βGeo</u> mice used in this study were originally generated at Bay Genomics and then back-crossed for several generations to obtain line stability. An expression cassette encoding a β-galactosidase neomycin resistance fusion protein (βGeo) was inserted into intron 50 of *Kmt2d* on mouse chromosome 15 and is predicted to generate a truncated Kmt2d protein with peptide sequence corresponding to the first 50 exons of *Kmt2d* fused to βGeo, but lacking the SET domain and therefore methyltransferase activity^31^. For breeding of Kmt2d+/βGeo mice, matings were set up using a heterozygous male and WT female. For all analyses, littermates were used. For genotyping the following 4 primers were used: CCGAGTACCTGAAAGGCGAA, CTAATCAGGGCTAAGGGCGA, GTGTGGAACCGCATCATTGAG, GTCTTTGAGCACCAGAGGACA. For generation of mouse glia and human neurons co-cultures newborn pups from CD1 (ICR, Charles River) were used unless otherwise stated in the result and figure caption section.

### Mouse hippocampal cultures

To generate cultured mouse hippocampal neurons, brains were dissected from P0 mice and rinsed quickly with ice-cold Hank’s Balanced Buffered Saline (HBSS) before continued dissection of the hippocampus, digestion with papain, and then seeding onto matrigel-coated coverslips in MEM medium (ThermoFisher Scientific). P0 was defined as the day the pups were born. On day in vitro (DIV) 1, the medium was changed to neurobasal medium containing B27. On DIV3, the medium was exchanged for medium containing 2 μM AraC, and cells were infected with lentiviruses encoding reporters for live imaging (see viral constructs and virus generation). Live imaging or immunostaining was performed at DIV14–16, when the cells are considered mature.

### Confocal Imaging and Immunocytochemistry

Immunofluorescence staining was performed essentially as described ^43^. Briefly, cultured human neurons were fixed in 4% paraformaldehyde and 4% sucrose in PBS for 20 min at room temperature, washed three times with PBS, and incubated in 0.2% Triton X-100 in PBS for 10 min at room temperature. Cells were blocked in PBS containing 5% goat serum for 1 h at room temperature. Primary antibodies were applied overnight at 4°C, cells were washed in PBS for three times, and fluorescent-labelled secondary antibodies (Alexa, 1:1000) were applied for 2 h at room temperature. The following antibodies were used in immunocytochemistry experiments: <u>MAP2</u> (CPCA-MAP2, EnCor; 1:2000), <u>pan-synapsin</u> (made for this paper, #7739 (aa 2-30 of human Syn1); 1:1,000), <u>synapsin-1</u> (10.22; Synaptic Systems; 1:1000), <u>bassoon</u> (219E1, Synaptic Systems; 1:1000), <u>vGAT</u> (117G4, Synaptic Systems; 1:1000), <u>Gephyrin</u> (mAb7a, Synaptic Systems; 1:1000), <u>vGluT1</u> (AB5905, Millipore, 1:1000), <u>PSD95</u> (MA1-046, Invitrogen, 1:1000), <u>KMT2</u>D (ab231239, Abcam, 1:500), <u>UTX</u> (D3Q1I, Cell Signaling, 1:500), <u>Parvalbumin</u> (PV27a, Swant, 1:1000), <u>Somatostatin-28</u> (SY-160F7, Synaptic Systems; 1:1000), <u>NECAB1</u> (HPA023629, Millipore, 1:1000), <u>GFP</u> (A11122, Invitrogen, 1:2000), <u>GFAP</u> (CPCA-GFAP, EnCor; 1:5000), <u>UTX</u> (ABE409, Millipore; 1: 500), <u>HepaCAM</u> (18177-1-AP, Proteintech; 1:500), <u>Neuroligin-2</u> (5E6, Synaptic Systems; 1:1000), <u>CD49f</u> (14-0495-82, Thermo; 1:500), <u>CD44</u> (MA4400, Thermo, 1:500), <u>GAT1</u> (274 104, Thermo; 1:500), <u>GLAST</u> (PA5-72895, Thermo; 1:2000), <u>Mono-Methyl-Histone H3 (Lys4)</u> (D1A9, Cell Signaling; 1:1600). Images were taken using a Nikon Ti2 eclipse A1HD25 confocal microscope system, with a 20x or 60x objective at RT.

### Immunofluorescent labeling of hippocampus

For immunohistochemistry, P30 mice were perfused with phosphate-buffered saline (PBS) and then PBS containing 4% PFA. Brains were dissected and rinsed quickly in ice-cold PBS before soaking in 30% sucrose. Brain tissues were embedded in optimal cutting temperature compound (OCT, Tissue Plus, Fisher) with embedding mold and frozen on dry ice. Embedded brain samples were stored at −80°C until further use. To sample the hippocampus in horizontal planes, multiple dorsal-ventral (horizontal) cryosections with a thickness of 30 um were collected. Brain sections were sliced using a cryostat (Leica Microsystems, CM3050-S) and then blocked using a solution containing goat serum and Triton-X100. Immunostaining was performed on slices while free-floating. Individual slices were mounted manually onto superfrost slides (VWR) using a paint brush.

### Calcium, iGluSnFR and iGABASnFR imaging

Calcium imaging was performed as previously described ^46,72^. Briefly, after lentivirally mediated expression of GCaMP8m for 2 weeks for mouse neural cultures, 2 months for human excitatory neurons, or 4 months for human inhibitory neurons, coverslips were gently placed into a dark imaging chamber with an open bottom and then washed twice with a washing buffer before time-lapse imaging was performed in calcium-imaging buffer (as indicated in figure captions: 2 (low ion buffer) or 4 (high ion buffer) mM CaCl_2_ and 4 (low) or 8 (high) mM KCl in 129 mM NaCl, 25 mM HEPES, 30mM glucose, 1 mM MgCl_2_, 10 µM Glycine, pH adjusted to 7.2-7.4). Washing buffer: same as imaging buffer, except 2.5 (low) or 5 (high) mM KCl and 1 (low) or 2 (high) mM CaCl_2_.

After 1–2 min equilibration in a 37°C incubator, calcium-imaging was performed on an inverted epi-fluorescence microscope (Nikon EclipseTS2R with pre-warmed sample stage) with a 488 nm filter. GCaMP8m fluorescence was recorded for 2 min at a frame rate of 10Hz (or 20Hz for astrocytes) using NIS Elements 5.30.05. The ROI was set to 1192×1192 Mono 16-bit using a 20x objective. For each condition, up to 10 fields of view were imaged per coverslip and a minimal 2 coverslips were used for each biological batch. All images were acquired using the same exposure time. Time-lapse imaging was performed in areas containing confluent neuron populations with few overlapping cell somata. Similarly, human neurons transduced with F-TetO-iGluSnFR3.857.GPI were imaged in cell-culture media (to avoid over-excitation in purely excitatory cells) at 40Hz with otherwise identical settings as for calcium imaging. Mouse hippocampal neurons were transduced with either FUW-iGluSnFR3.857.GPI or FUW-iGABA-SnFR2 and imaged at 40Hz in high ion buffer to trigger activity in these mixed excitatory and inhibitory cultures. Videos with multiple neurons were recorded. Imaging files were analyzed using MATLAB through custom-made code. Neurons or astrocytes were selected and then each were analyzed to generate statistics summary: frequency and mean/standard deviation of amplitudes for each neuron and synchronous spikes (dependent on the sigma threshold we set, 1.5). For each replicate, the data was combined and graphed using GraphPad Prism: Amplitude, Frequency, Synchronous Amplitude, and Synchronous Frequency.

### Hippocampal slice electrophysiology

Mice were anesthetized with isoflurane followed by ketamine/xylazine, then rapid cardiac perfusion was performed using ice-cold sucrose artificial cerebrospinal fluid (ACSF) containing the following (in mM): 85 NaCl, 2.5 KCl, 1.25 NaH_2_PO_4_, 25 NaHCO_3_, 25 glucose, 75 sucrose, 0.5 CaCl_2_, and 4 MgCl_2_, equilibrated with 95% O_2_/5% CO_2_ before mice were decapitated. Brains were rapidly removed and placed in the same ice-cold sucrose ACSF as described previously ^73^. Horizontal hippocampal slices (250 μm thick) were prepared using a Leica VT1000S or VT1200 S vibratome and transferred to a heated (28-32°C) holding chamber containing the same sucrose ACSF, which was gradually exchanged for regular ACSF containing the following (in mM): 125 NaCl, 2.5 KCl, 1.25 NaH_2_PO_4_, 25 NaHCO_3_, 25 glucose, 2 CaCl_2_, and 1 MgCl_2_, equilibrated with 95% O_2_/5% CO_2_ at room temperature. After incubation in regular ACSF for at least one hour, slices were transferred to a recording chamber and perfused continuously with oxygenated regular ACSF at a flow rate of 2 ml/min.

Excitatory and inhibitory postsynaptic currents (EPSCs and IPSCs) were recorded from pyramidal neurons in the CA1 subregion of the hippocampus. Stimulating electrodes were placed in the stratum radiatum to stimulate Schaffer collaterals from the CA3 neurons. EPSCs and IPSCs were isolated by switching the holding voltage at-50 mV and at 0 mV, which are close to the reversal potentials for IPSCs and EPSCs ^74^. Input-output (I-O) curves were derived by plotting current amplitudes corresponding to each incremental stimulating intensity (10 - 20 μA increments). Stimulation intensity was adjusted to evoke EPSCs with amplitudes of 300-400 pA in individual neurons, and then IPSCs were recorded from the same neurons to estimate the inhibitory to excitatory (I/E) ratio.

### Viral constructs and virus generation

The following lentiviral constructs were used: <u>FUW-TetO-Ngn2-T2A-puromycin</u> (TetO promoter drives expression of full-length mouse Ngn2 and of puromycin via the cleavage-peptide sequence T2A)^43^, <u>FUW-</u> <u>rtTA</u>^43^, <u>FUW-GFP::Cre</u> to express Cre-recombinase or <u>GFP::ΔCre</u> for the wild-type control^45^. <u>FUW-GCaMP8m</u> for Ca²^+^ live imaging (modified for this paper from original version pGP-CMV-jGCaMP8m that was a gift from GENIE Project^75^ (Addgene plasmid # 162372; http://n2t.net/addgene:162372; RRID:Addgene_162372<u>), F-</u> <u>GFAP-GCaMP8m</u> for Ca²^+^ live imaging in astrocytes (modified for this paper form original plasmid^76^), <u>F-TetO-</u> <u>iGluSnFR3</u> (iGluSnFR live imaging in human induced neurons; modified for this paper from original version FSW_iGluSnFR3.857.GPI that was a gift from Ronald Hart (Addgene plasmid # 205986; http://n2t.net/addgene:205986; RRID:Addgene_205986)), <u>FUW-iGluSnFR3</u> (iGluSnFR live imaging in mouse neurons; modified for this paper from original version FSW_iGluSnFR3.857.GPI), <u>FUW-iGABA-SnFR2</u> (iGABASnFR live imaging in mouse neurons; modified for this paper from original version pGP-CAG-iGABASnFR2n-WPRE-bGH-polyA was a gift from GENIE Project (Addgene plasmid # 218869; http://n2t.net/addgene:218869; RRID:Addgene_218869)^76^), <u>FUW-EGFP</u> (for sparse transfection of human neurons)^77^, F-TetO-Ascl1-T2A-puromycin^56^, F-TetO-Dlx2-hygromycin^56^, <u>lentiCRISPR v2</u> (used to insert specific gRNA sequences)^78^.

Lentiviruses were produced as described ^43^ in HEK293T cells (ATCC, VA) by co-transfection with three helper plasmids (pRSV-REV^79^, pMDLg/pRRE^79^ and vesicular stomatitis virus G protein expression vector (pMD2.G that was a gift from Didier Trono (Addgene plasmid # 12259; http://n2t.net/addgene:12259; RRID:Addgene_12259)) with 12 µg of lentiviral vector DNA and 6 µg of each of the helper plasmid DNA per 75 cm^2^ culture area) using calcium phosphate. Lentiviruses were harvested in the medium 48 h after transfection, aliquoted, and stored at-80°C.

### Human induced neurons (iN cells) and Glia co-culture

Experiments were performed as follows. 1019 iPSCs (ATCC-DYS0100 Human Induced Pluripotent Stem (IPS) Cells) were maintained as feeder-free cells in Stemflex medium (Thermo). Human neuron generation has been described previously ^43^. Briefly, targeted human iPSCs were treated with Accutase (Innovative Cell Technologies) and plated as dissociated cells in 24-well plates (1 x 10^4^ cells/well) on day-1. Cells were plated on matrigel (BD Biosciences)-coated coverslips in Stemflex containing 2 mM thiazovivin (Bio Vision). At the same time point, lentiviruses were prepared as described above and added to each well (14 µl/well of 24-well plate) in a total volume of 500 µl. Mainly two different types of lentiviruses were used for co-infection: the lentiviruses used for human neuron induction as described, and lentiviruses expressing either Cre-recombinase (to create a null allele) under control of the ubiquitin promoter or an inactive mutated ΔCre-recombinase for the wild-type control. On day 0, the culture medium was replaced with N2/DMEM/F12/NEAA (Invitrogen) containing human BDNF (10 ng/ml, PeproTech), human NT-3 (10 ng/ml, PeproTech), and mouse Laminin-1 (0.2 µg/ml, Thermo). Doxycycline (2 µg/ml, Clontech) was added on day 0 to induce TetO gene expression and retained in the medium until the end of the experiment. On day 1, a 24 hr puromycin selection (1 µg/ml) (and hygromycin 200 µg/ml) selection for inhibitory iNs) period was started. On day 2, mouse glia cells to excitatory iNs (to inhibitory iNs on day 9) were added in neurobasal medium supplemented with B27/Glutamax (Thermo) containing BDNF, NT3 and Laminin-1; Ara-C (2 µM, Sigma) was added to the medium to inhibit glial proliferation. Mouse glial cells were cultured from the forebrain of new-born wild-type CD1 mice. Briefly, new-born mouse forebrain homogenates were digested with papain and EDTA for 15 min, cells were dissociated by harsh trituration to avoid growing of neurons, and plated onto T75 flasks in DMEM supplemented with 10% FBS (Corning) and Pen/Strep (Thermo). Upon reaching confluence, glial cells were trypsinized and replated at lower density a total of two times to remove potential trace amounts of mouse neurons before the glia cell cultures were used for co-culture experiment with human neurons. After day 2 (excitatory iNs) or 9 days (inhibitory iNs), 50% of the medium in each well was exchanged every 2 days. FBS (5%) was added to the culture medium on day 10 to support astrocyte viability, and human neurons were assayed after at least 2 (excitatory) or 4 (inhibitory) months.

### Generation of KMT2D-Conditional and KDM6A-Conditional Knockout iPSCs

To create a human KMT2D conditional knockout model, CRISPR-Cas9 plasmids were designed to sequentially (first upstream then downstream) target the regions flanking exon 6 of KMT2D in otherwise wild-type male iPSCs (1019 line), which upon deletion would result in the loss of 163 bp of coding DNA and thereby induce a shift in the open reading frame resulting in heterozygous loss and haploinsufficiency as observed in patients. Single-guide RNAs (sgRNAs) were selected to specifically target sequences upstream (GAGAAGGACCATGGGTATAC) and downstream (GTGTGAACTTTGTGAGCACC) of exon 6 (cloned into lentiCRISPR v2 ^78^), and the following donor templates containing loxP sites were constructed to facilitate homology-directed repair: CAGGGGATTGCATAGGTGAAGGGCAGGGCTGGTGGGCTTCTGAGAGTCAGGTTGGAATGAGAAGGACCATGGGTAATAACTTCGTATAGCATACATTATACGAAGTTATTACTGGGGCAGAGCTAGTGCCTGAACTGTTTGTCTGGGGAGAGCTGGCTGACACTGAGGCTCTTTTTTCACCCTG and TACAGAGCACTGATCAAAGTATAGCATTATTTACTGTTGGTCTGCCCCACTAAAATAGGTGTGAACTTTGTGAGCATAACTTCGTATAGCATACATTATACGAAGTTATACCAGGGTTTTGCCTGTCTCATTTATCATTTTGTCTCCAGTGCCTAGAACAGTAGCTGGCAGACGTTGGGCAGGC.

After nucleofection (4D-Nucleofector, Lonza) with the lentiCRISPR v2 plasmid and donor single stranded DNA templates (IDT), cells were subjected to antibiotic selection using puromycin (1 µg/ml) for 48 hours, beginning 1 day post-transfection. Surviving iPSC colonies were allowed to expand for an additional 10–14 days and were subsequently picked and screened by PCR for successful loxP integration. Targeted clones were identified by PCR using specific primers designed to detect the upstream and downstream loxP insertions. The following primer pairs were used for verification: The upstream loxP site was confirmed using the oligos AGATCAGGACTTCTCACCCAGATATGCAGC (forward) and GCTTCTGAGAGTCAGGTTGGAATGAGAAGGA (reverse) while the downstream loxP site was confirmed using the oligos GGCACTCCATGGGCACAGAATGAAC (forward) and ATCTGGGTGAGAAGTCCTGATCTTTGGTCA (reverse).

For the upstream LoxP site, a correctly targeted KMT2D allele yielded a PCR product of 333 bp, while untargeted wild-type alleles produced a 299 bp band. For the downstream LoxP site, a correctly targeted KMT2D allele yielded a PCR product of 302 bp, while untargeted wild-type alleles produced a 268 bp band. Colonies showing the expected band sizes were expanded, and correct targeting was confirmed by sequencing. Following verification, a selected clone was cryopreserved and further validated by differentiation into neurons, where conditional knockout could be induced by Cre recombinase to ensure functional loxP integration and recombination.

For KDM6A / UTX, the sequential targeting strategy focused on exon 7. Similar to the KMT2D approach, CRISPR-Cas9 was used to flank exon 7 (121 bp) with loxP sites. The following sgRNAs were designed and ligated into the lentiCRISPR v2 plasmid to target the sequences surrounding exon 7: CACTATGAGTATTGTCATAG (upstream), TAATTTTTATCCCACGCAGT (downstream). A donor template with loxP sites was used for homology-directed repair in otherwise wild-type male iPSCs (1019 line), yielding a hemizygous mutation / deletion, as observed in male patients. iPSCs were nulceofected with the CRISPR-Cas9 plasmid and the UTX donor template, followed by puromycin selection (1 µg/ml) starting 1 day after nucleofection. Following selection, individual colonies were expanded and screened by PCR using primers specific for the upstream and downstream loxP insertions: The upstream loxP site was confirmed using the oligos ATGAACTTGGACACAGGCTATTTTAAAAGGCAC (forward) and GGGTTATTTTCTTGAGACAGCTGAGAATGGAG (reverse) while the downstream loxP site was confirmed using the oligos CCTGCAAGACCAGATATTGTGAACAGCCT (forward) and AATCAACCCAATCTGCTTTTGAAAAGACACCT (reverse). For the upstream loxP site, a correctly targeted UTX allele yielded a PCR product of 393 bp, while untargeted wild-type alleles produced a 259 bp band. For the downstream loxP site, a correctly targeted UTX allele yielded a PCR product of 460 bp, while untargeted wild-type alleles produced a 426 bp band. Colonies showing the expected band sizes were expanded, and correct targeting was confirmed by sequencing. Following verification, a selected clone was then used in differentiation protocols to derive neurons and assess UTX function post-knockout induction via Cre recombinase.

### Axon and Dendrite Morphology Analysis

To selectively visualize and analyze axon morphology, human neurons were sparsely transfected with FUW-EGFP (using the calcium phosphate method) at DIV19 to label individual neurons and their axonal processes distinctly. This sparse GFP labeling allowed for clear visualization of axon structures, enabling measurements of axonal length and branching. Neurons were fixed at DIV21 with 4% paraformaldehyde (PFA) in phosphate-buffered saline (PBS) for 10 minutes at room temperature and subsequently washed three times with PBS. Cells were simultaneously permeabilized and blocked with 5% goat serum + 0.1% Triton X-100 in PBS for 1 hour. For dendrite analysis, neurons were incubated overnight at 4°C with primary antibodies against MAP2. After primary antibody incubation, cells were washed with PBS and treated with Alexa Fluor-conjugated secondary antibodies for 1 hour at room temperature in the dark. Coverslips were mounted using Fluoromount-G (Invitrogen, 00-4958-02). Cells were imaged using a 10x objective and analyzed for parameters including axon length, axon branching, dendrite length, dendritic branching, and soma size. Axon length was measured manually using the NIS Elements Line Tool to trace the GFP+/MAP2-axon projection extending from the soma to its distal end. Axon branching was determined by visually counting the primary branches extending from the main axon process. Dendrites labeled by MAP2 were analyzed using MetaMorph software (Molecular Devices) and assessed for total dendrite length (summing the lengths of all dendritic branches extending from the soma, number of processes (counting the primary dendritic processes emerging from the soma), number of branches (counting secondary and tertiary branches of dendrites), and soma size (measuring the cross-sectional area of the soma). All measurements were performed across 2-4 coverslips for each condition, each sampled from three biological replicates. Results were expressed as mean ± SEM, and statistical significance was determined using the Student’s t-test for comparing two groups.

### RNA sequencing

RNA was extracted from DIV14 hippocampal cultures established from KS1-mutant and wild-type P0 mouse littermates, with cultures derived from three independent litters. RNA-seq library preparation and sequencing was performed at the Notre Dame Genomics and Bioinformatics Core Facility. Quality of RNA samples was confirmed using High Sensitivity RNA ScreenTape (Agilent) before Illumina libraries were prepared from 200 ng of RNA using the NEBNext Ultra II Directional RNA Library Prep kit with Sample Purification Beads (E7765S) and using the NEBNext Poly(A) mRNA Magnetic Isolation Module (E7490S). Libraries were confirmed using the Fragment Analyzer and quantified by qPCR prior to pooling and sequencing on an Illumina NovaSeq using 101 bp paired-end reads. On average, 1 NextSeq P4 XLEAP flow cell generates 1.5-1.8 Billion clusters/reads, for an average of 125 Million raw reads per sample across both lanes. Bulk RNA seq was performed for a total of 3 biological replicates per condition. The same library was sequenced twice as technical replication and to ensure adequate depth.

### Analysis of RNA-seq data

Raw reads were processed with fastp v0.23.4 to only use reads > 60 bp and phred score Q > 30 ^80^. Reads were mapped to GRCm39 using hisat2 v2.10 ^81^. The R package DESeq2 v1.40.2 was used to analyze differential expression ^82^. Y chromosome genes were removed from the analysis to avoid calling DE on the basis of sex. The collapseReplicates() function from DESeq2 was used to collapse technical replicates. DESeq2 internally performs library size and count normalization. Variance partitioning ^83^ and PCA were used to confirm a lack of batch effects. DEseq2 calculated Benjamin-Hochberg adjusted p-values with a default alpha of 0.1. Adjusted p-value ≤0.05 and LFC cutoff of ±0.5 (corresponding to an approximately 40% change in expression) were used to determine significantly differentially expressed genes. The org.Mm.eg.db v3.17.0 annotation package was used for gene set enrichment analysis by clusterProfiler v4.8.3 ^84^. LFC values from the DESeq2 results were used to rank genes and set sizes were limited to a min and max of 100 and 500 respectively. Semantic similarity clustering of GO terms was performed using GOSemSim v2.34.0 to reduce the redundancy of select plotted GO terms using Wang’s method ^85^. All scripts are available at: https://github.com/nmcadam2.

Bulk RNAseq does not allow for individual investigation of constituent cell types as is possible in single cell analyses. However, the observed expression values from RNAseq are the additive result of the number of cells from each cell type weighted by the expression patterns of those cell types. As a result, we can estimate cell type mathematically if we have the observed data (bulk RNAseq expression) and the expression pattern of each cell type individually using deconvolution via BayesPrism ^86^. This is a baysian model that treats the single-cell derived data as a prior and estimates the joint posterior distribution of cell type fraction in each bulk sample. We performed standard BayesPrism analysis on the bulk RNA sequencing results from mouse-mouse samples in this study, using single-cell data from Allen Brain Map Mouse Whole Cortex and Hippocampus SMART-seq of 77k total cells to define expression patterns for each cell type ^87^. Hippocampal regions were merged as we were not interested in individual regions. The single-cell data included data for glutamatergic, GABAergic, and non-neuronal (glia) cells. Individual glia subtypes were not identified in the input data. The top 5000 markers were used for both glutamatergic and non-neuronal cells, but only 142 GABAergic markers could be used due to the limitation of information in the input single cell data. This analysis resulted in the predicted composition of cell types in each sample (Suppl. Fig. 12d).

The BayePrism results were evaluated with cluster correlation analyses as part of standard practice. The cluster correlation analysis of individual regions showed no internal structure, supporting our ability to merge all hippocampus regions. Cluster correlation of cell types showed the expected closer alignment of expression in neurons versus non-neuronal cells.

### Data presentation and statistics

Part of Figures 3b, 5f, and 7 were created with BioRender.com. All data shown are individual values and means ± SEMs; Number of measured cells and/or independent experiments is indicated inside each bar or mentioned in the figure. All statistical analyses were performed using the unpaired two-tailed Student’s t-test or the Mann-Whitney U test, one-way ANOVA, two-way ANOVA repeated measurements or Kolmogorov-Smirnov test for cumulative distributions, comparing the test sample to the control sample examined in the same experiments. No power analysis was performed to predetermine sample sizes. Our sample sizes are similar to those generally employed in the field. Sample allocation was random. Statistical significances were calculated using Graphpad Prism.

## ACKNOWLEDGEMENTS

This work was supported by funds from the University of Notre Dame, the Brain foundation to C.P. and NIH 1R01CA255064-01A1 to Siyuan Zhang. We thank the undergraduate summer students Aaron Davis, Andrew Mauriello and Jeremiah Lopez for helping with specific experimental analyses.

## AUTHOR CONTRIBUTIONS

Conceptualization: A.W., J.K., C.P.

Data curation: A.W., J.K., S.F., N.M., T.N., S.S., A.C., C.P.

Formal analysis: A.W., J.K., S.F., A.W., N.M., S.M., T.N., A.W., A.N., C.S., S.S.

Funding acquisition: A.W., J.K., C.P.

Investigation: A.W., J.K., S.F., A.W., N.M., S.M., T.N., A.W., J.S., A.N., C.L., S.N., C.M., I.W.-S., C.S., K.H., J.B.-P., S.S., C.P.

Methodology: A.W., J.K., N.M., T.N., C.L., S.N., C.M., S.S., C.P.

Project administration: C.P. Resources: K.H., A.C., C.P.

Software: N.M., C.M., C.S., S.S.

Supervision: A.C., C.P.

Validation: A.W., J.K., N.M., S.S., C.P.

Visualization: A.W., J.K., N.M., T.N., C.P. Writing - original draft: A.W., J.K., C.P.

Writing - review & editing: A.W., J.K., S.F., A.W., N.M., S.M., T.N., A.W., J.S., A.N., C.L., S.N., C.M., I.W.-S., C.S., K.H., J.B.-P., S.S., K.H., A.C., C.P.

## SUPPLEMENTARY FIGURES AND SUPPLEMENTARY FIGURE LEGENDS

**Suppl. Figure 1:**
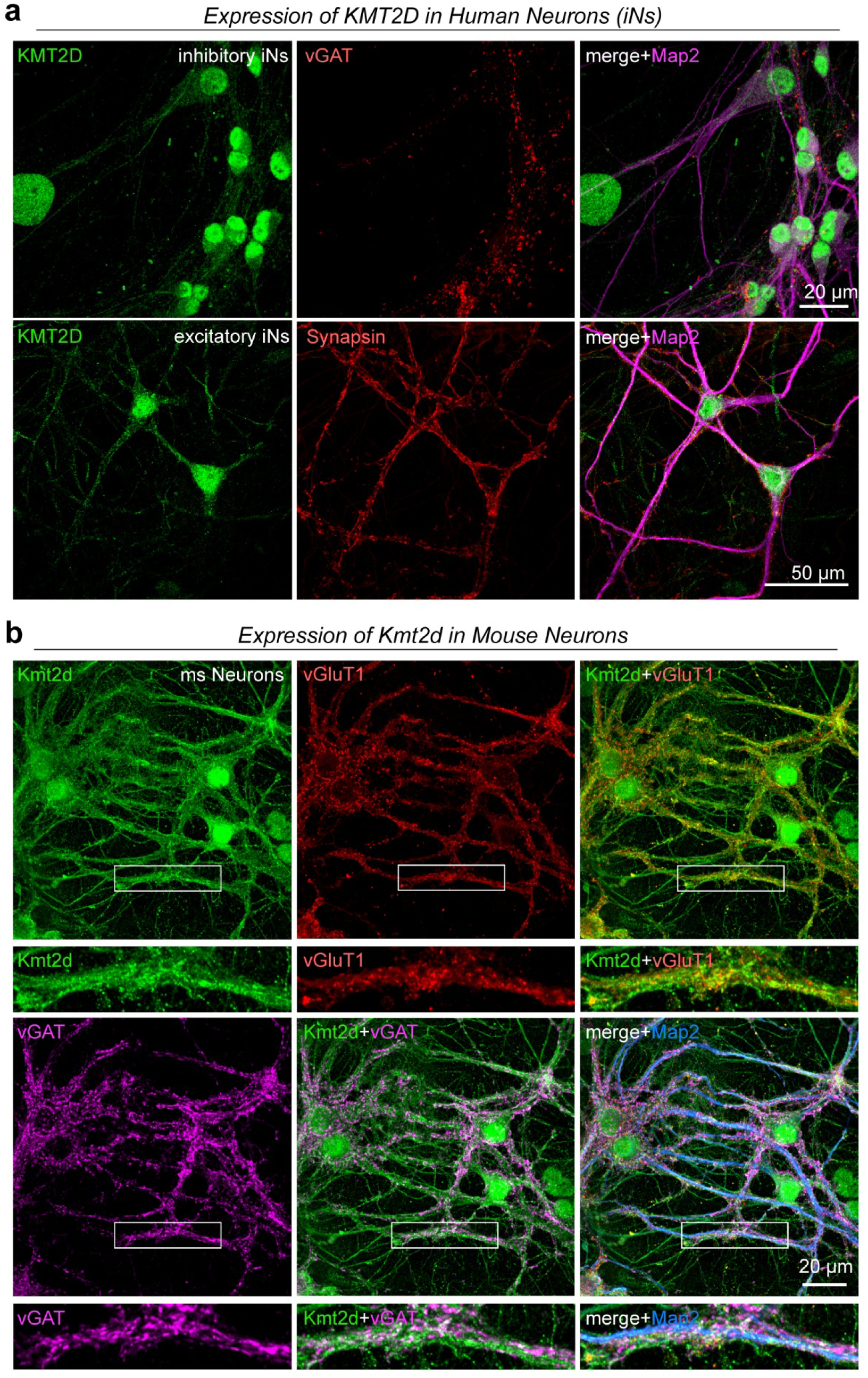
Expression of KMT2D in wild-type human and mouse neurons, Related to Figure 1 a, Representative micrographs showing KMT2D (green) expression in inhibitory wild-type human induced neurons (iNs; upper panels, generated by overexpression of Ascl1 and Dlx2 in iPSCs) and excitatory iNs (lower panels, generated by overexpression of Ngn2). KMT2D localization is shown in relation to the presynaptic inhibitory vesicle marker vGAT (red), the pan-presynaptic vesicle marker Synapsin-1 (red), and the postsynaptic dendritic marker MAP2 (magenta). Co-cultured wild-type mouse glia also show robust Kmt2d expression. b, Micrographs of Kmt2d expression (green) relative to the presynaptic excitatory vesicle marker vGluT1 (red), the presynaptic inhibitory vesicle marker vGAT (magenta) and the postsynaptic dendritic marker MAP2 (blue) in cultured wild-type mouse hippocampal neurons.

**Suppl. Figure 2:**
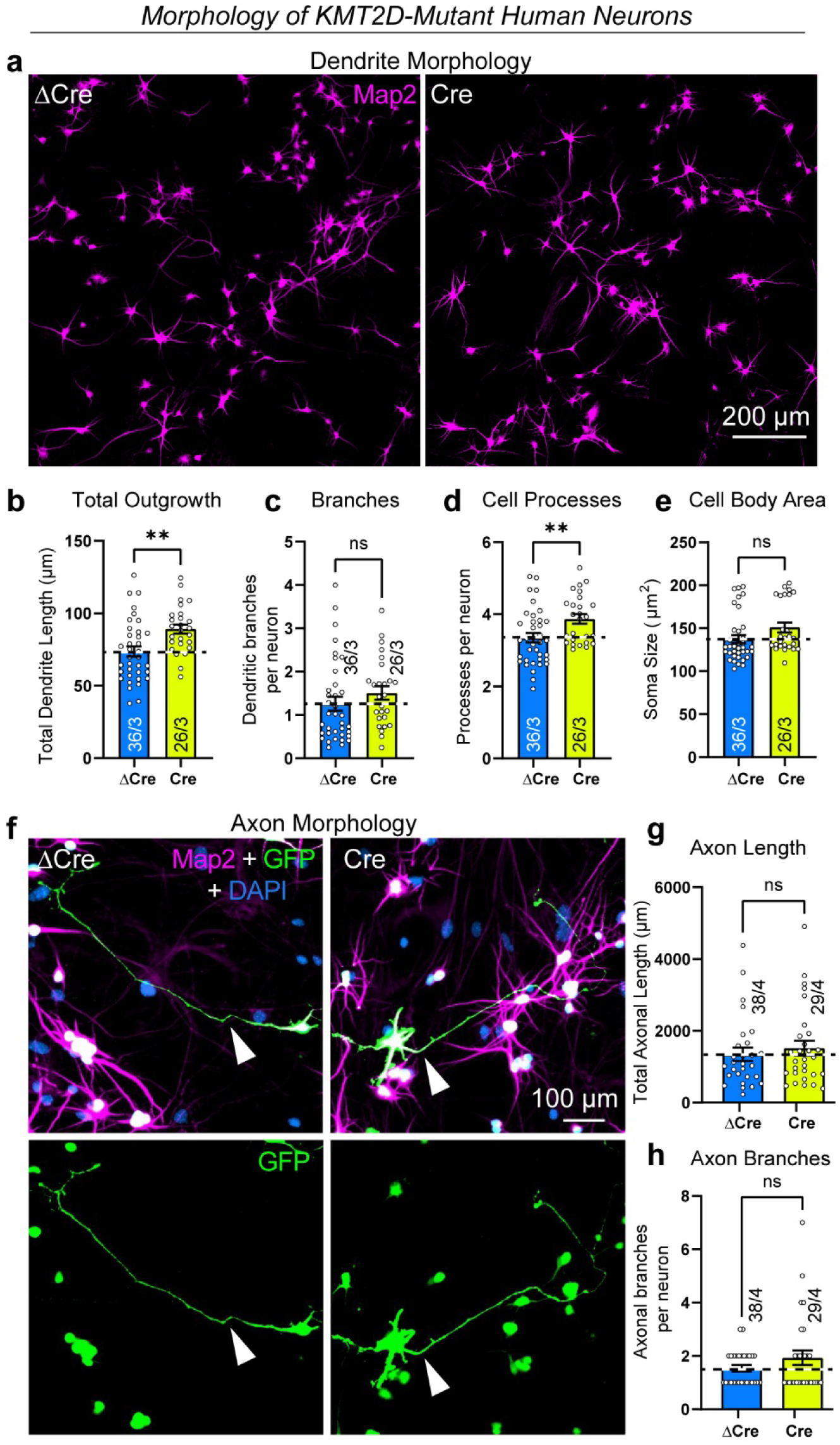
Dendrite length and complexity are enhanced in *KMT2D*-mutant human neurons (KS1) while no differences were found for axon morphology, Related to Figure 1 a, Representative images of dendritic morphology in human excitatory neurons stained with MAP2 (magenta) from ΔCre (*KMT2D* wild-type) and Cre (*KMT2D* mutant) cultures at 21 days. b-e, Quantification of dendritic outgrowth, processes, branches, and soma size in ΔCre and Cre neurons. B, Total dendrite length shows a significant increase in Cre mutant neurons compared to ΔCre neurons. c, No significant differences in dendritic branches per neuron between ΔCre and Cre neurons. d, Number of cell processes per neuron is significantly increased in Cre mutant neurons. e, Cell body area (soma size) also shows no significant differences between ΔCre and Cre neurons. f, Representative images of axonal morphology, with dendrites stained by MAP2 (magenta), nuclei stained with DAPI (blue), and axons visualized via sparse EGFP-transfection (green) in ΔCre and Cre neurons (with green nuclei due to ΔCre-GFP and Cre-GFP fusion proteins). g, No significant differences in total axon length between ΔCre and Cre neurons. h, No significant differences in the number of axon branches per neuron between ΔCre and Cre neurons. Data are individual values (each representing one image in b-e or one cell in g and h) and means ± SEM; number of cells or images analyzed and number of independent cell cultures are shown in graphs; statistical significance was assessed by two-tailed unpaired Student’s t-test (**=p<0.01).

**Suppl. Figure 3:**
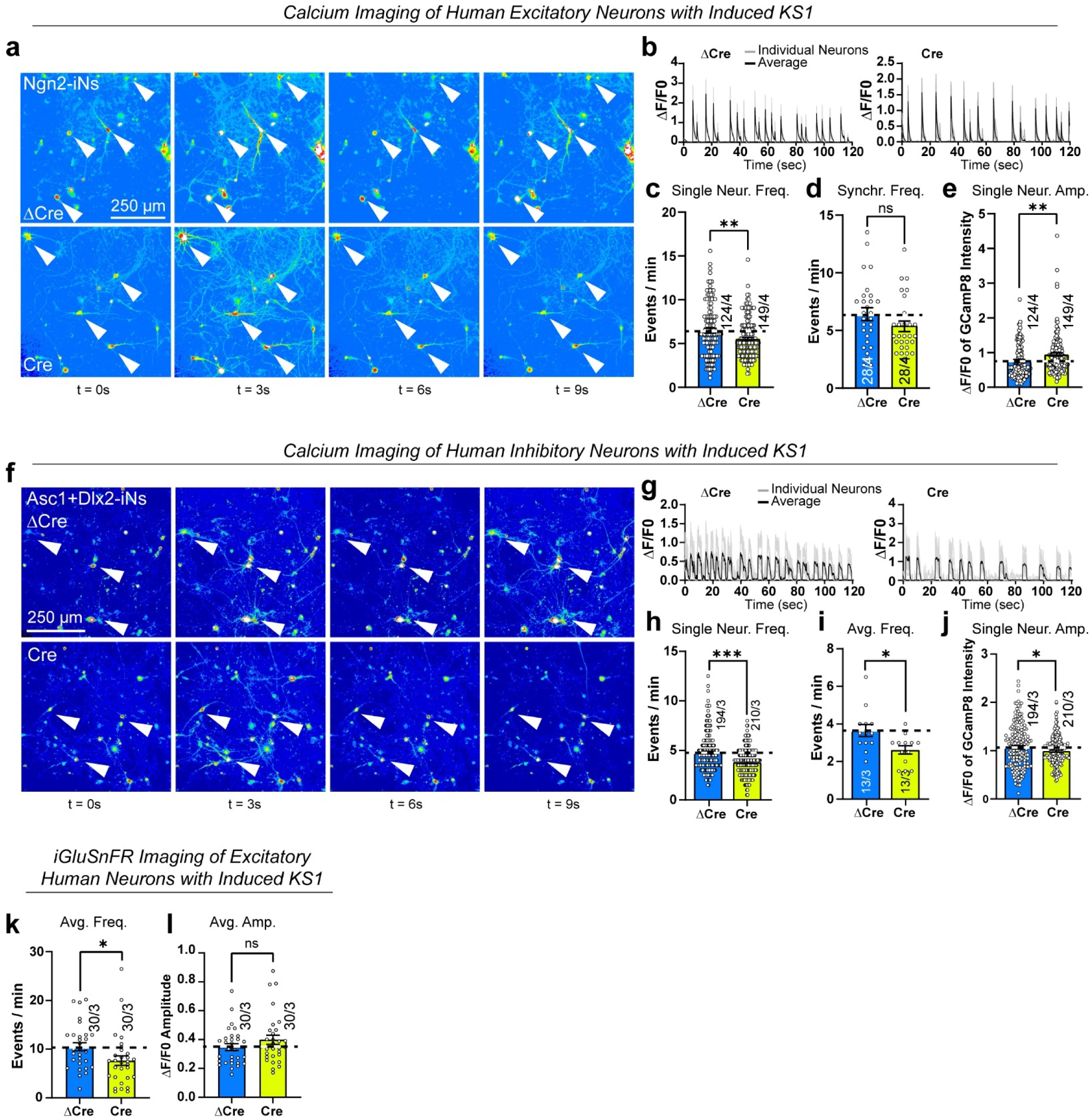
Reduced calcium transients and excitatory synaptic transmission frequency in the human neuron model of Kabuki syndrome type 1 (KS1), Related to Fig. 1 a, Representative calcium imaging micrographs of cultured wild-type (ΔCre) and mutant (Cre) excitatory human neurons overexpressing GCaMP8, recorded at 10Hz. Neuronal activity was induced by 2.5 mM KCl and 2 mM CaCl_2_. Warmer colors indicate higher fluorescence intensity. Arrows highlight active neurons displaying calcium transients. b, Representative traces of GCamp8m transients showing fluorescence intensity changes (ΔF/F_0_) over time. Mutant neurons show decreased event frequency. c-e, Summary graphs of individual neuron firing frequency (c), synchronous firing frequency per view field (d), and signal intensities in single neurons (e). f, Representative calcium imaging snapshots showing changes in calcium activity in wild-type (ΔCre) and mutant (Cre) inhibitory human induced neurons over time (t = 0, 3, 6, and 9 seconds) analogous to A. g, Calcium event traces of GCaMP8m intensity (ΔF/F_0_) over 120 seconds in ΔCre and Cre neurons, recorded at 10Hz. Each spike represents a calcium event, with Cre neurons showing decreased frequency and heightened amplitude compared to ΔCre neurons. h-i, Quantification of calcium event frequency and amplitude in ΔCre and Cre neurons. Single neurons and average frequency show significant decrease in Cre compared to ΔCre neurons. k-l, Summary graphs show average values of event frequency and amplitude per cell per field of view. Data from individual cells are shown in Fig. 1o-p. Data are individual values (each representing one video in d, i, k, l; and one cell in c, e, h and j) and means ± SEM; number of cells or videos analyzed and number of independent cell cultures are shown in graphs; statistical significance was assessed by unpaired Student’s t-test (*, p<0.05; ** p<0.01; ***=p<0.001).

**Suppl. Figure 4:**
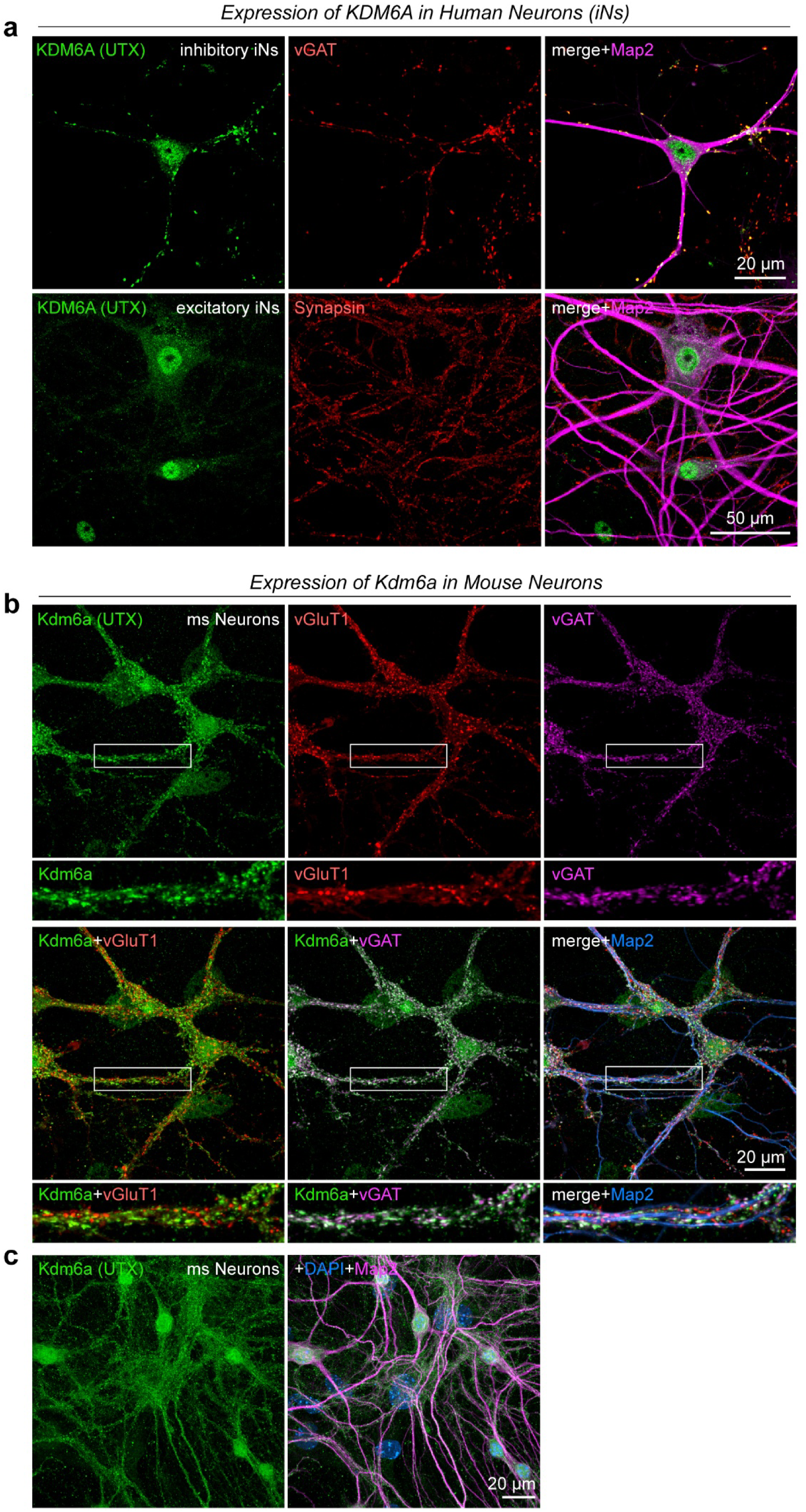
Expression of KDM6A (UTX) in wild-type human and mouse neurons, Related to Figure 2 a, Representative micrographs showing KDM6A (green) expression in inhibitory wild-type human iNs (upper panels) and excitatory iNs (lower panels). KDM6A localization is shown in relation to the presynaptic inhibitory vesicle marker vGAT (red), the pan-presynaptic vesicle marker Synapsin-1 (red), and the postsynaptic dendritic marker MAP2 (magenta). Co-cultured wild-type mouse glia also show robust Kmt2d expression. b, Micrographs of Kdm6a expression (green) relative to the presynaptic excitatory vesicle marker vGluT1 (red), the presynaptic inhibitory vesicle marker vGAT (magenta) and the postsynaptic dendritic marker MAP2 (blue) in cultured wild-type mouse hippocampal neurons. c, Expression of Kdm6a revealed by a different antibody (as in a and b) in wild-type mouse hippocampal neurons.

**Suppl. Figure 5:**
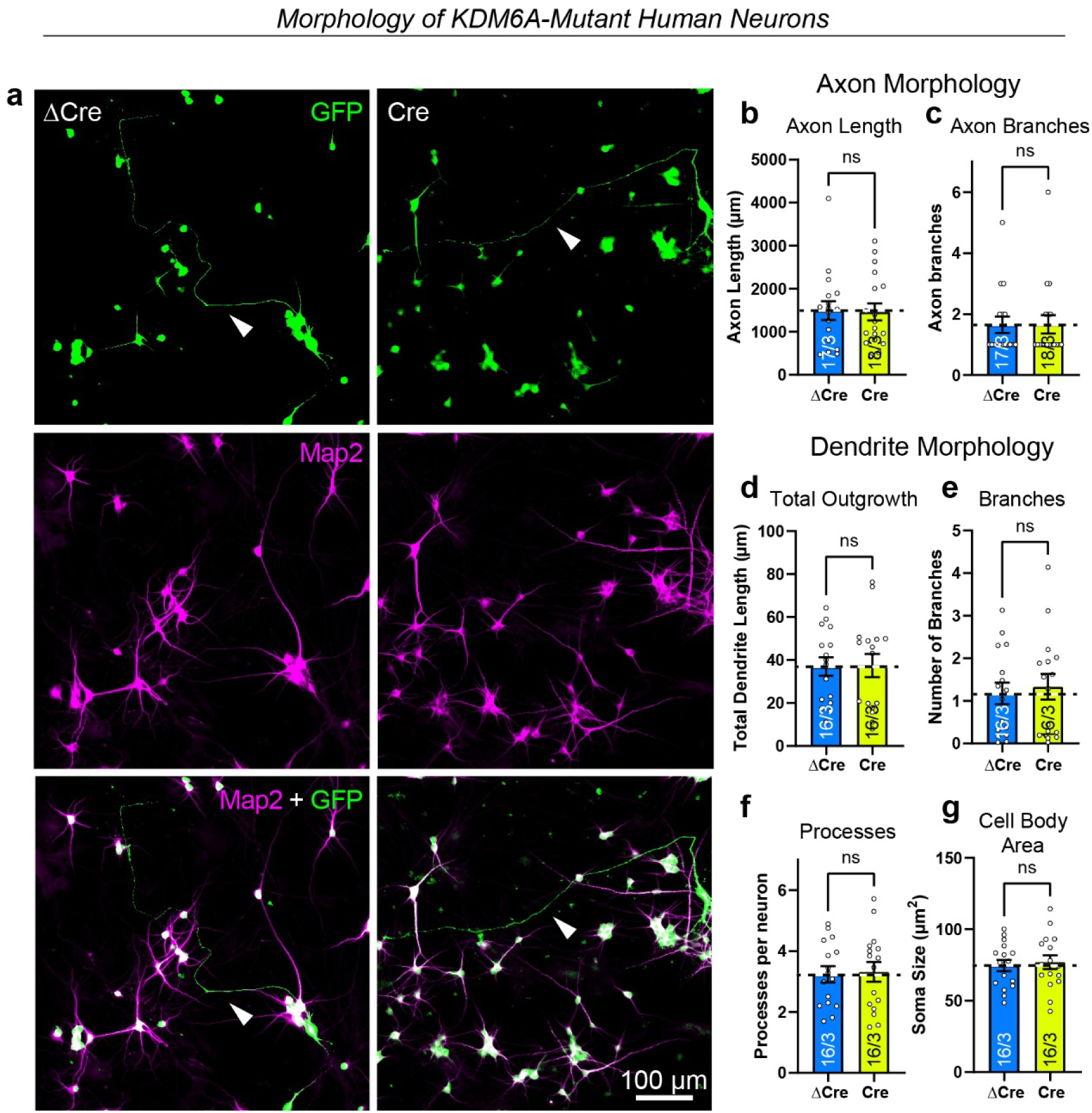
Conditional mutation of *KDM6A* (KS2) in human excitatory neurons leads to no differences in dendrite or axon morphology, Related to Figure 2 a, Representative images of axon and dendrite morphology in ΔCre (wild-type) and Cre (*KDM6A* mutant) Ngn2-derived neurons. Composite images (bottom row) display sparse overexpression of EGFP (green, marking axons) and MAP2 (magenta, marking dendrites), with arrows indicating axonal structures. EGFP images (middle row) show axonal structures (green), while MAP2 images (bottom row) depict dendritic structures (magenta) in ΔCre and Cre neurons. b-g, Quantification of axonal and dendritic properties in ΔCre and Cre neurons. Data are individual values (each representing one cell in b and c or one image in d-g) and means ± SEM; number of cells or images analyzed and number of independent cell cultures are shown in graphs; statistical significance was assessed by unpaired Student’s t-test.

**Suppl. Figure 6:**
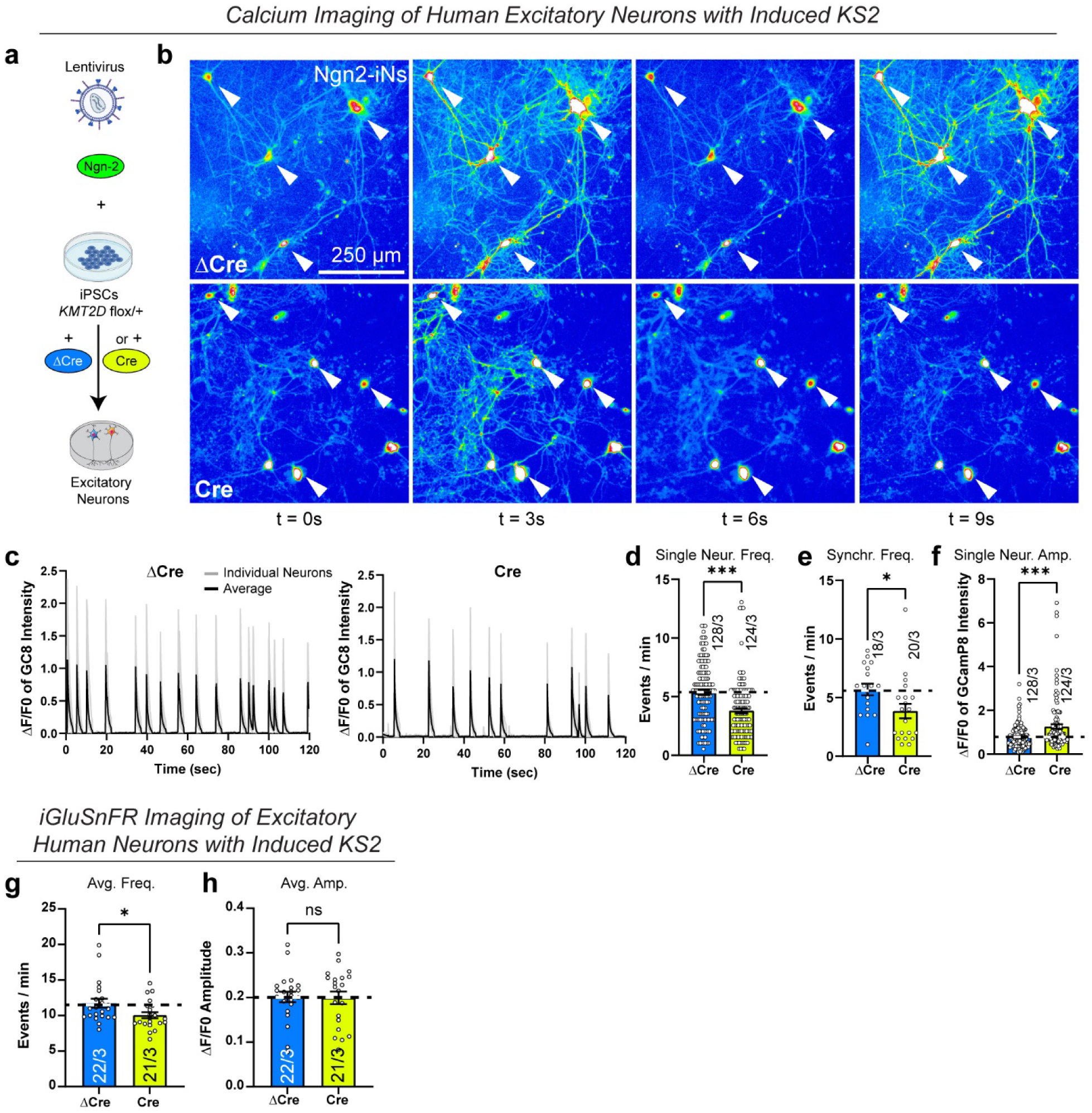
Reduced calcium transients and excitatory synaptic transmission frequency in the human neuron model of Kabuki syndrome type 2 (KS2), Related to Fig. 2 a and b, Differentiation protocol used for generating excitatory iNs (overexpression of transcription factor Ngn2) for subsequent calcium imaging at 2 months. Representative calcium imaging micrographs of wild-type (ΔCre) and mutant (Cre) human iNs overexpressing GCaMP8m. Neuronal activity was induced by 8 mM KCl and 4 mM CaCl_2_. Warmer colors indicate higher fluorescence intensity. Arrows point to active neurons displaying calcium transients. c, Representative traces of GCaMP8m transients showing fluorescence intensity changes (ΔF/F_0_) over time. Mutant neurons show decreased event frequency. d-f, Summary graphs of individual neuron firing frequency (d), synchronous firing frequency per view field (e), and signal intensities in single neurons (f). g-h, Summary graphs show average values of event frequency and amplitude per cell per field of view. Data from individual cells are shown in Fig. 2k-l. Data are presented as individual values (each representing one video in e, g, h; and one cell in d and f) and means ± SEM; number of cells or videos analyzed and number of independent cell cultures are shown in graphs; statistical significance was assessed by unpaired Student’s t-test (*, p<0.05; *** p<0.001).

**Suppl. Figure 7:**
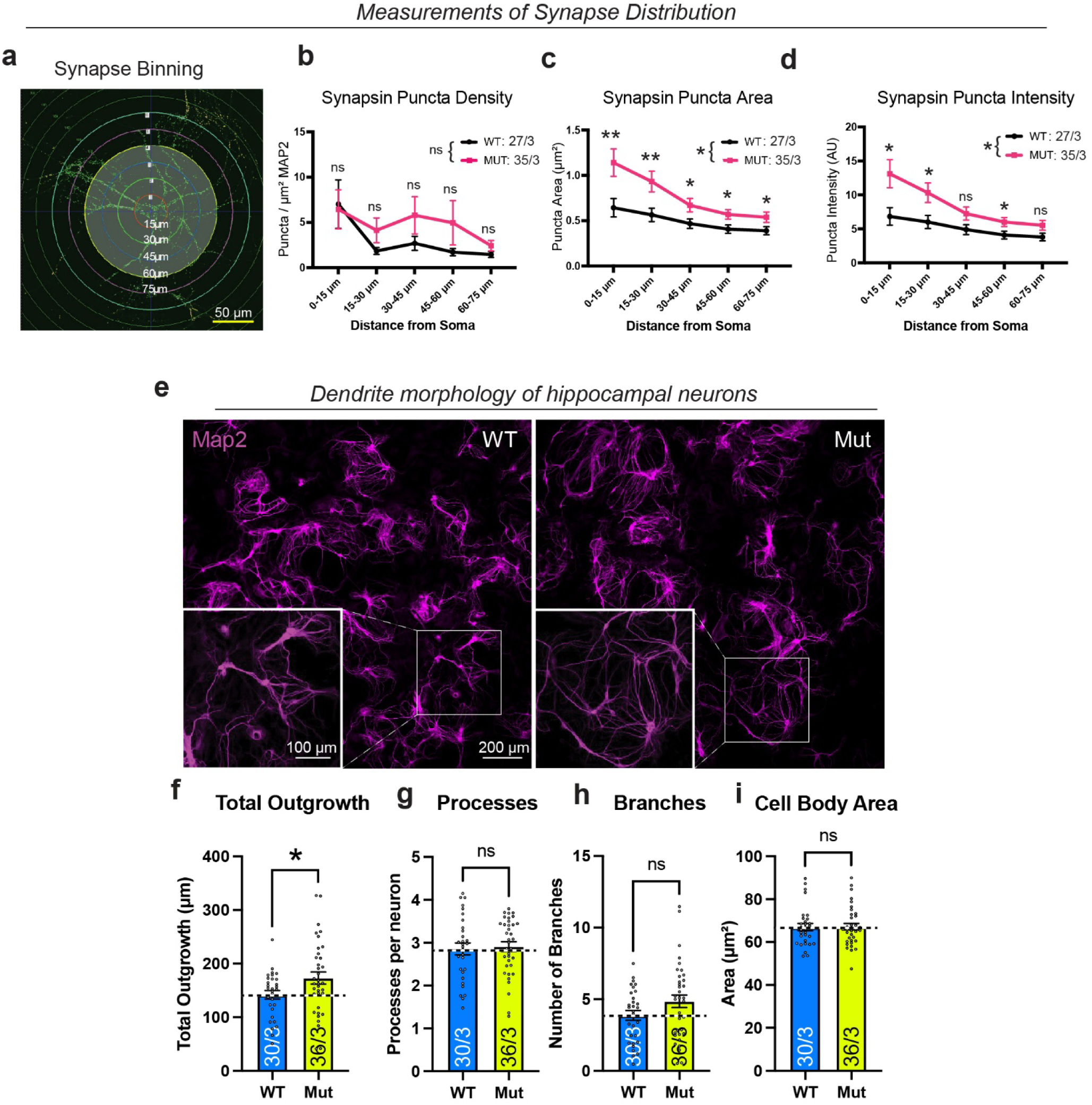
*Kmt2d* mutation (KS1) leads to changes in overall synapse distribution and increased dendritic arbor in cultured hippocampal neurons, Related to Figure 3 a, Schematic representation of synapse binning used to measure synapse distribution at different distances from the soma in mouse hippocampal neurons. b-d, Summary graphs of synaptic puncta density, area, and intensity as a function of distance from the cell soma. Mutant neurons show a greater increase in Synapsin-1 puncta metrics, particularly at distances closest to the soma. e, Dendritic outgrowth is enhanced in Kmt2d-mutant neurons. Immunocytochemistry images show Map2-labeled dendrites in wild-type (WT, left) and mutant (Mut, right) hippocampal neurons cultured for 2 weeks. Insets provide zoomed-in views of dendritic structures. f-i, Quantification of dendritic morphology in WT and Mut neurons: f, Mutant neurons show a significant increase in total dendritic outgrowth compared to WT neurons. g, No significant difference in the number of processes per neuron (ns = not significant). h, There was no significant difference in dendritic branching between WT and Mut neurons. i, No significant difference was observed in the cell body area between WT and Mut neurons. Data are presented as individual values (each representing one image) and/or means ± SEM in b-c; number of images analyzed and number of independent cell cultures are shown in graphs statistical; significance in b-d and f-i was assessed by two-tailed unpaired Student’s t-test (*, p<0.05; **, p<0.01) and additionally in b-d by two-way ANOVA repeated-measurements (*, p<0.05).

**Suppl. Figure 8.**
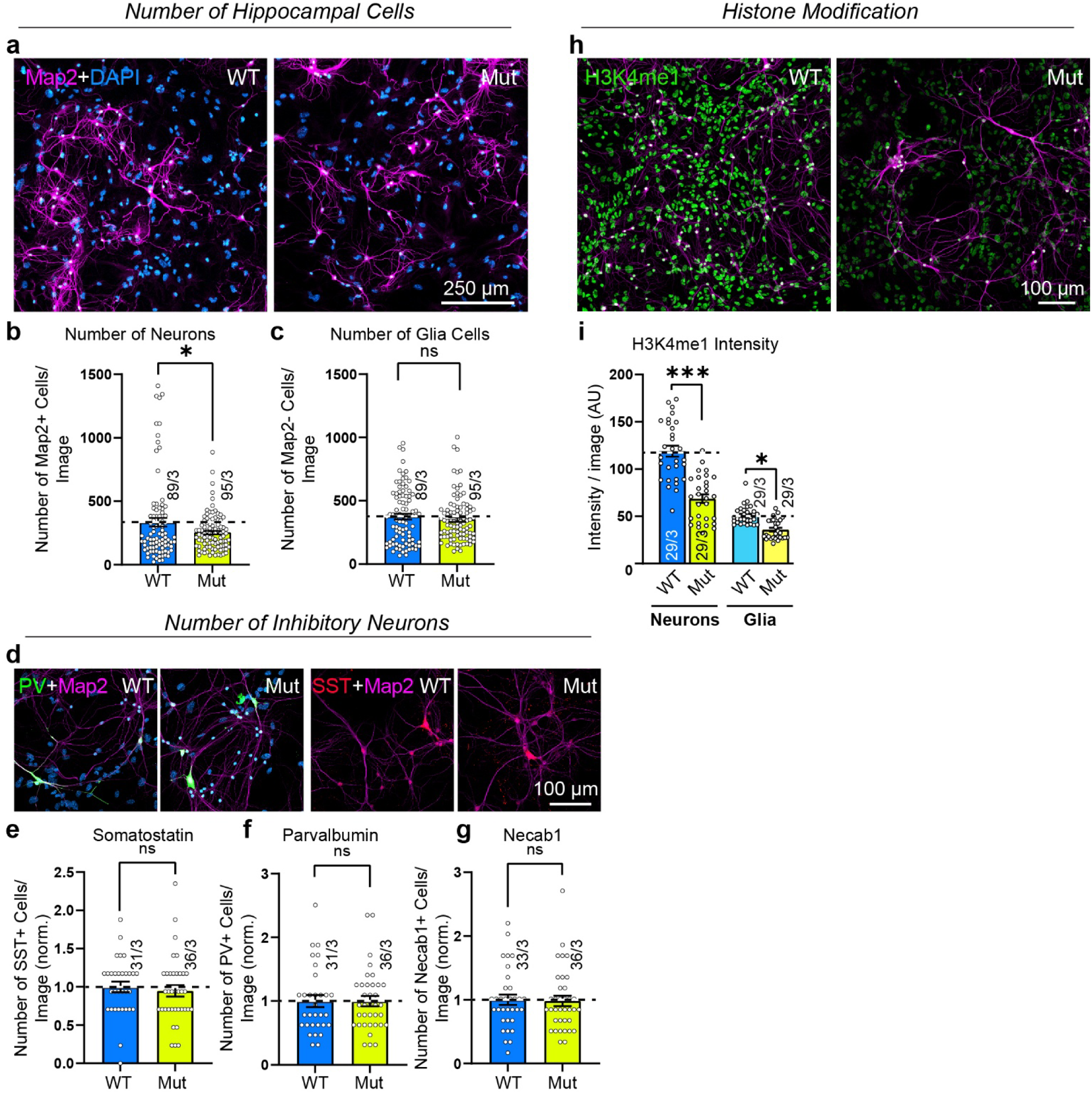
*Kmt2d* mutation in mice (KS1) leads to decreased number of overall hippocampal neurons but no change in the number of inhibitory neurons, Related to Figure 3 a-c, Representative micrographs of WT and Mut hippocampal cells. Neuronal numbers were assessed by Map2-positive somata. Map2-negative cells were considered glia cells. d-g, Representative micrographs showing WT and Mut hippocampal cells stained for inhibitory neuronal markers Somatostatin (SST) and Parvalbumin (PV). Summary graphs quantify the relative (to WT) number of positive cells, including the inhibitory neuronal marker Necab1. h and i, *Kmt2d*-mutant neurons (Mut) neurons and glia exhibit decreased Histone 3 lysine 4 mono-methylation. Data are presented as individual values (each representing one image) and means ± SEM; number of images analyzed and number of independent cell cultures are shown in graphs; statistical significance was assessed by unpaired Student’s t-test (*, p<0.05; *** p<0.001).

**Suppl. Figure 9:**
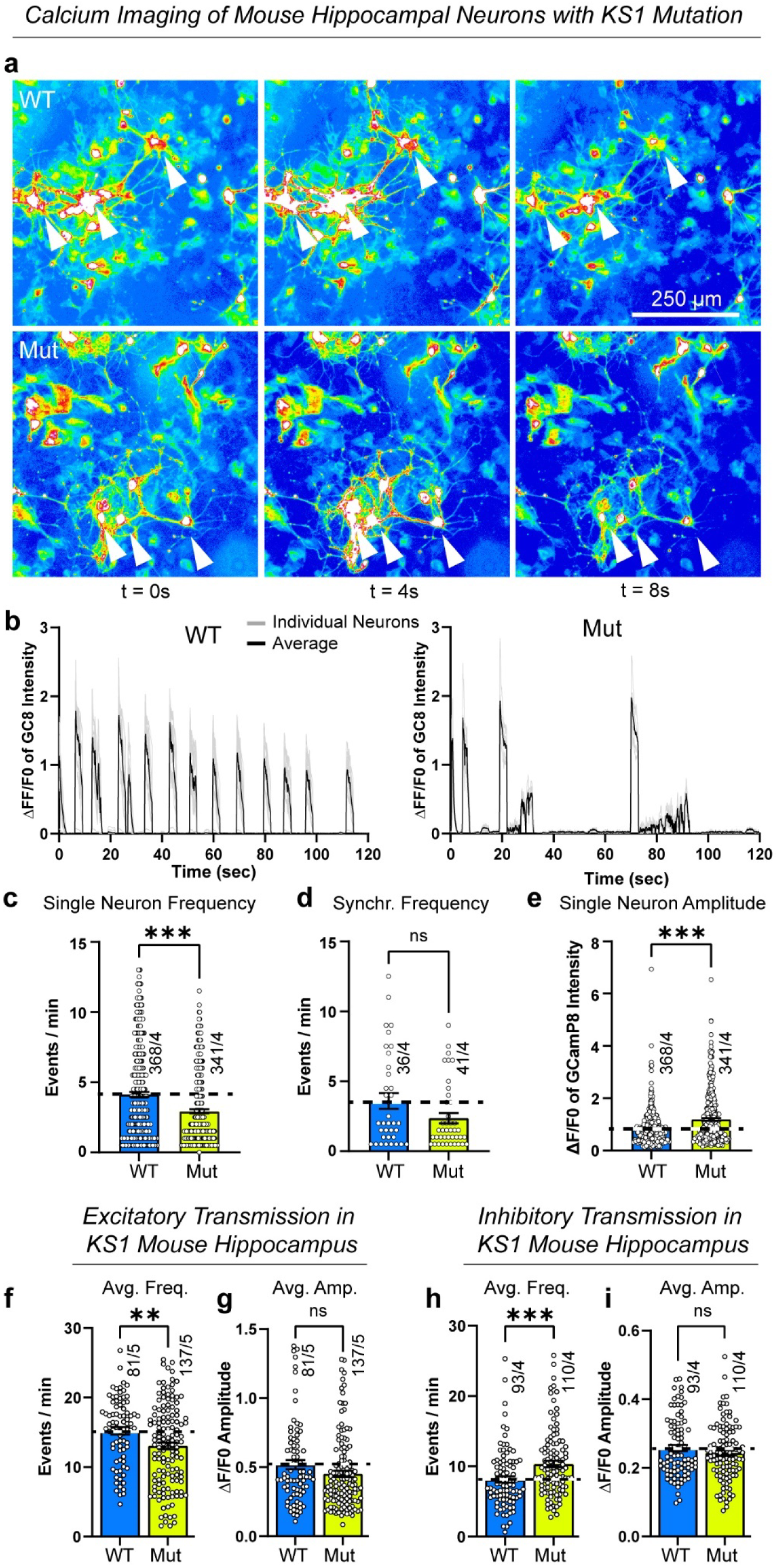
*Kmt2d* mutation (KS1) in mice leads to decreased neuronal calcium activity, Related to Figure 4 a, Representative calcium imaging micrographs of cultured wild-type (WT) and mutant (Mut) mouse hippocampal neurons overexpressing GCaMP8m. Network activity was induced with 2.5 mM KCl and 2 mM CaCl_2_. Warmer colors indicate higher fluorescence intensity. Arrows highlight active neurons showing calcium transients. b, Representative traces of GCaMPm8 transients. Fluorescence intensity changes (ΔF/F0) over time reveal a more clustered pattern and higher signal intensities in Mut neurons. c-e, Summary graphs showing individual neuron firing frequency (c), synchronous firing frequency per view field (d), and signal intensities in single neurons (e). f-g, Summary graphs show average values of iGluSnFR-event frequency and amplitude per cell per field of view. Data from individual cells are shown in Fig. 4d-e. h-i, Summary graphs show average values of iGABASnFR-event frequency and amplitude per cell per field of view. Data from individual cells are shown in Fig. 4i-j. Data are presented as individual values (each representing one movie) and means ± SEM; number of cells or videos analyzed and number of independent cell cultures are shown in graphs; statistical significance was assessed by unpaired Student’s t-test (**=p<0.01, *** p<0.001).

**Suppl. Figure 10:**
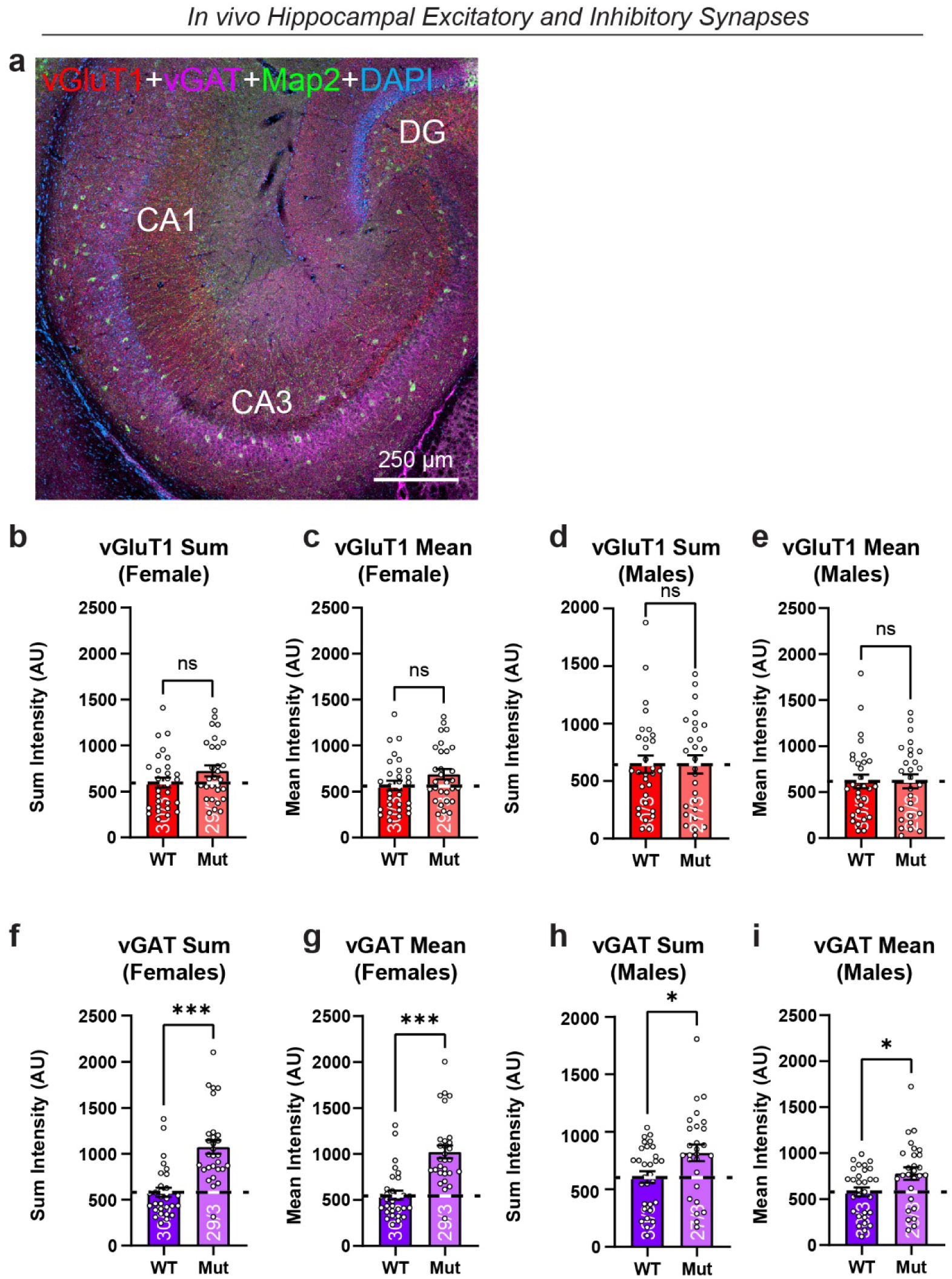
Increased number of inhibitory synapses in male and female *Kmt2d*-mutant (KS1) mice *in vivo*, Related to Figure 5 a, Overview image of hippocampal formation in a young adult (P30) WT female mouse. Section were stained for Map2 (green), vGluT1 (red), vGAT (magenta), and DAPI (blue). b-i, Summary graphs for vGAT and vGluT1 staining showing mean instensity (intensity of all pixels/number of pixels) and the sum intensity of all pixels with the field of view. Images analyzed correspond to those shown in Figure 5a. Data here are identical to Figure 5b-e but are separated into male and female mice. Data are presented as individual values (each representing one image) and means ± SEM; number of images analyzed and number of independent animals are shown in graphs; statistical significance was assessed by unpaired Student’s t-test (*, p<0.05; *** p<0.001).

**Suppl. Figure 11:**
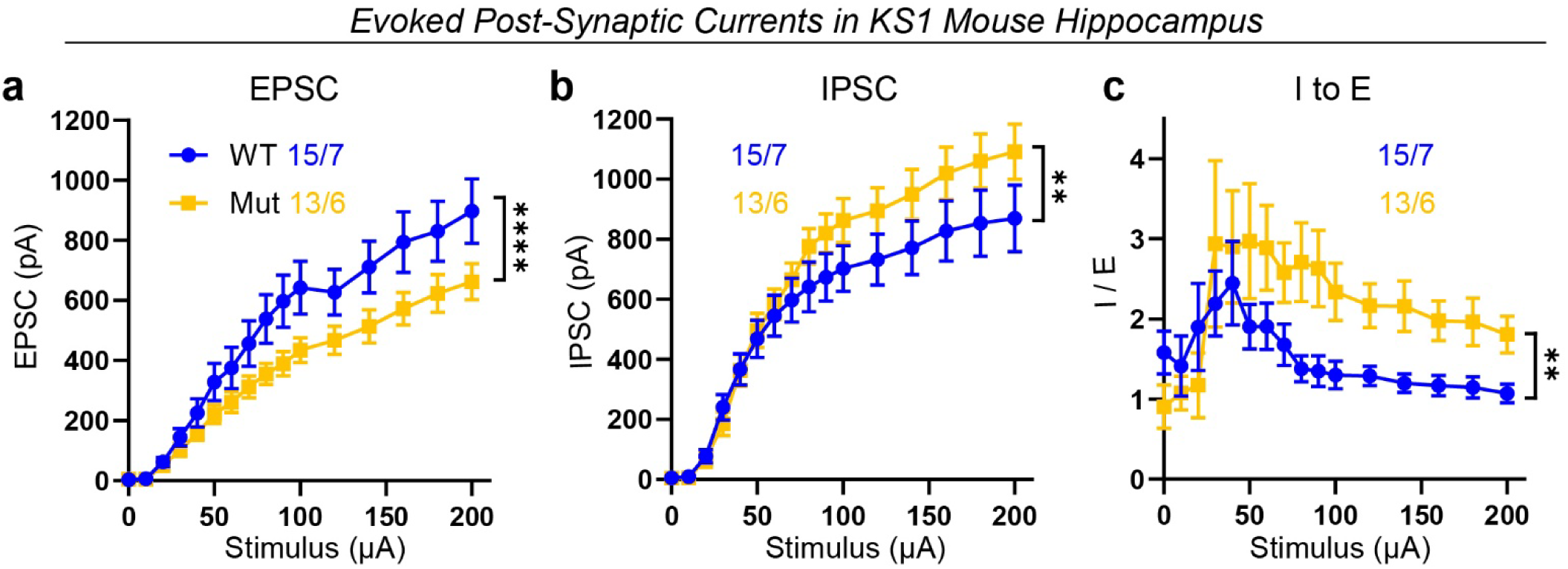
Decreased evoked excitatory and increased evoked inhibitory synaptic transmission in *Kmt2d*-mutant (KS1) *ex vivo*, Related to Figure 5 a and b, Input-Output (I-O) curves of excitatory post-synaptic currents (EPSCs (a)) or inhibitory post-synaptic currents (IPSCs (b)) following incremental stimulation steps of 20 μA in WT and *Kmt2d*-mutant mice. WT: 15 cells from 7 mice; Mut: 13 cells from 6 mice were recorded. c, Ratio of IPSCs (a) and EPSCs (b) recorded from the same cell. Data are individual values (representing one cell) and means ± SEM; number of cells analyzed and number of independent animals are shown in graphs; statistical significance was assessed by two-way ANOVA repeated-measurements (**, p<0.01; ****, p<0.0001).

**Suppl. Figure 12:**
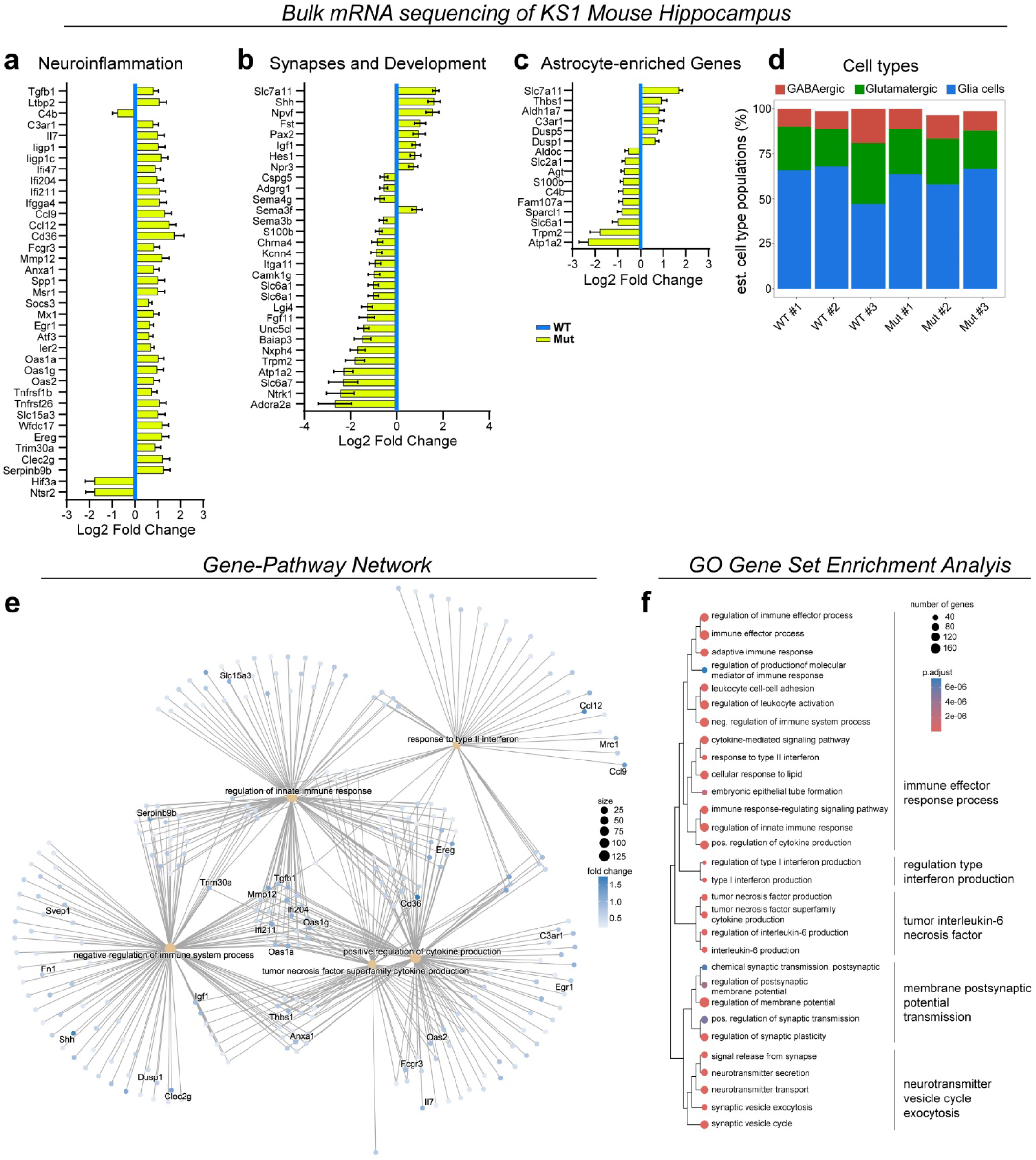
*Kmt2d* haploinsufficiency (KS1) in the mouse hippocampus triggers neuroinflammation and expression changes in glia enriched genes, Related to Figure 6 a-c, Mutant mice select differentially expressed genes (yellow) vs. wild type (blue). Graph shows log2 fold change and p adj as error bars. d, Cell type and gene expression deconvolution of sequenced hippocampal cells using BayesPrism. e, Gene ontology (GO) term to gene graph showing relative expression of leading edge genes for select pathways. Size: number of genes in the leading edge. f, Dot plot showing normalized enrichment scores (NES) for select GO terms. Clustering is based on semantic similarity of GO terms. Size: number of genes. N=3 biological replicates from 3 experiments.

**Suppl. Figure 13:**
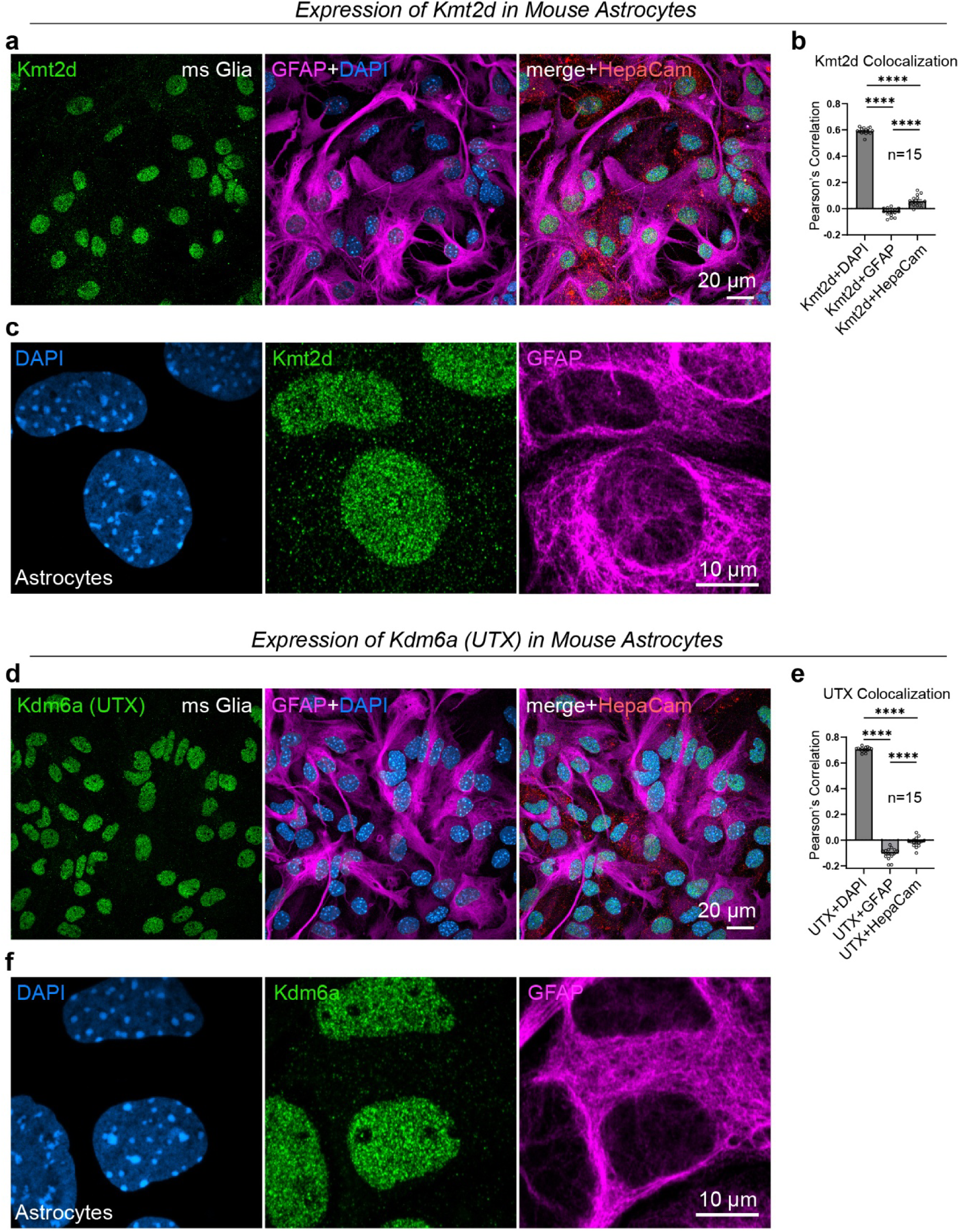
Expression of Kmt2d and Kdm6a in wild-type mouse glia cell, Related to Figure 6 a and c, Low (a) and high (a) magnification images wild-type mouse glia cells showing expression of Kmt2d (green) in relation to astrocyte markers GFAP (magenta) and HepaCAM (red). Nuclei are visualized by DAPI (blue). b, Summary graph of Pearson’s correlations for colocalization of Kmt2d with mentioned markers. d and f, Low (d) and high (f) magnification images wild-type mouse glia cells showing expression of Kdm6a (Utx) (green) in relation to astrocyte markers GFAP (magenta) and HepaCAM (red). Nuclei are visualized by DAPI (blue). e, Summary graph of Pearson’s correlations for colocalization of Kdm6a with mentioned markers. Data are presented as individual values (each representing one image) from 1 experiment and means ± SEM; statistical significance was assessed by unpaired Student’s t-test (**** p<0.0001).

**Suppl. Figure 14:**
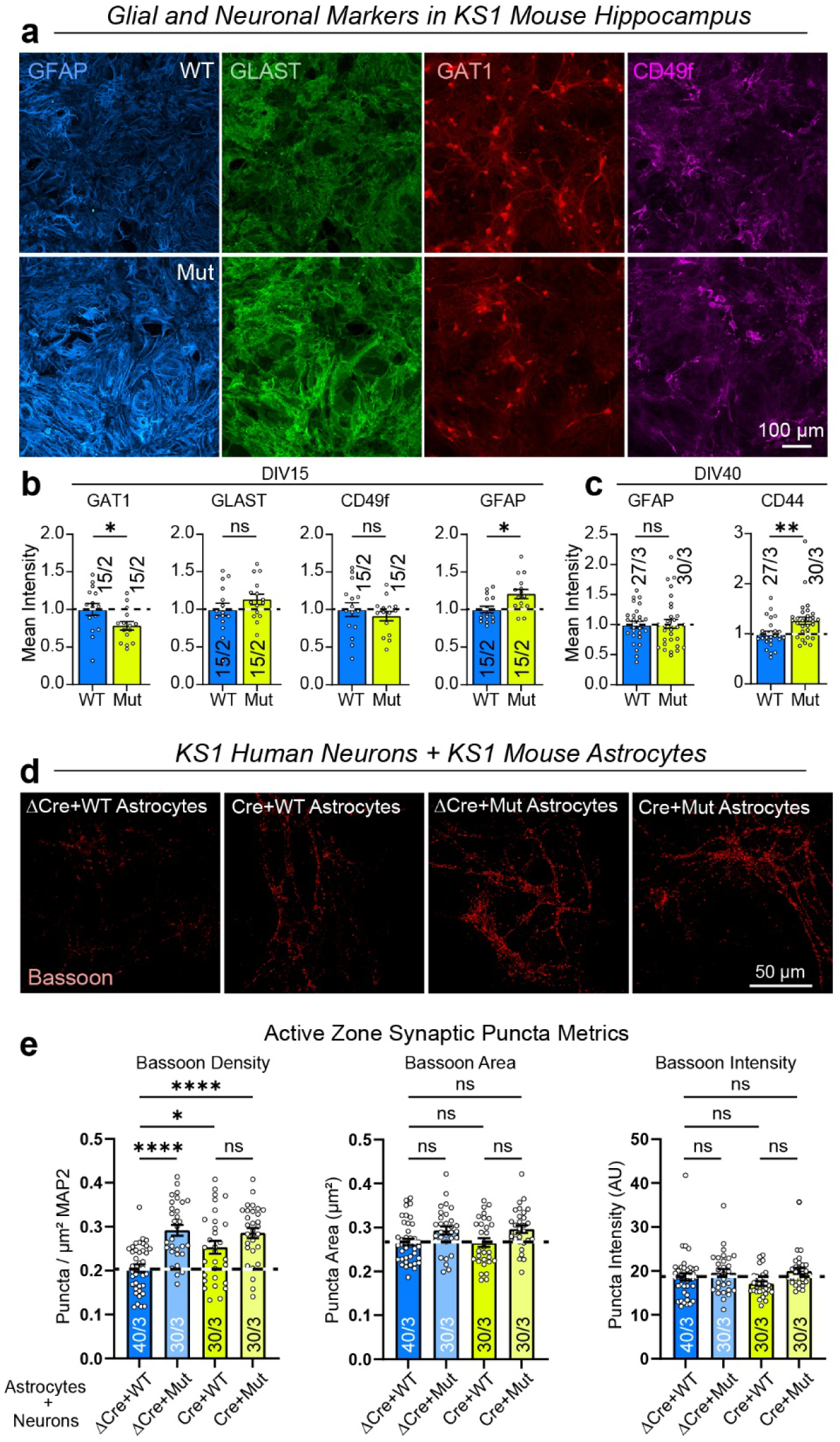
*Kmt2d* haploinsufficiency (KS1) in the mouse hippocampus transiently triggers reactive astrocytes and reduced GAT-1 expression, driving increased inhibitory synapse formation in human neurons, Related to Figure 6 a-c, Representative micrographs showing WT and Mut hippocampal cells immune-labeled for Glial Fibrillary Acidic Protein (GFAP, blue), Excitatory Amino Acid Transporter 1 (GLAST, green), GABA transporter type 1 encoded by *Slc6a1* (GAT1, red), and Integrin alpha 6 (CD49f, magenta). Summary graphs show their intensity levels at different times. Additionally, levels of CD44 (a marker for inflammation in astrocytes; increased) are shown. d, Co-culture of human wild-type (ΔCre) or *KMT2D*-mutant induced inhibitory neurons (Cre) with wild-type (WT) or *Kmt2d*-mutant (Mut) mouse astrocytes (same view field as Fig. 6h). e, KS1 mutations in human neurons or mouse astrocytes are sufficient to increase synaptic puncta metrics (revealed by pre-synaptic marker Bassoon) in inhibitory human neurons. Data shown are presented as individual values (each representing one image) with indicated mean ± SEM; number of cells or images analyzed and number of independent cell cultures are shown in graphs. Statistical significance was assessed by unpaired Student’s t test for summary graphs in b and c; and one way ANOVA and Tukey’s post hoc test for multiple comparisons in E (*, p<0.05; **, p<0.01; **** p<0.0001).

